# AtRBOHC/RHD2 vesicular delivery to the apical plasma membrane domain during root hair development

**DOI:** 10.1101/2021.09.13.460100

**Authors:** Lenka Kuběnová, Michaela Tichá, Jozef Šamaj, Miroslav Ovečka

**Affiliations:** Department of Cell Biology, Centre of the Region Haná for Biotechnological and Agricultural Research, Faculty of Science, Palacký University Olomouc, Šlechtitelů 27, 783 71 Olomouc, Czech Republic

**Keywords:** advanced microscopy, *Arabidopsis thaliana*, RBOHC, RHD2, root hair, tip growth, vesicular trafficking, plasma membrane

## Abstract

Arabidopsis root hairs develop as long tubular extensions from the rootward pole of trichoblasts and exert polarized tip growth. The establishment and maintenance of root hair polarity is a complex process involving the local apical production of reactive oxygen species (ROS) generated by NADPH oxidase RESPIRATORY BURST OXIDASE HOMOLOG PROTEIN C/ROOT HAIR DEFECTIVE 2 (AtRBOHC/RHD2). It has been shown that loss-of-function *rhd2* mutants have short root hairs that are unable to elongate by tip growth, and this phenotype was fully complemented by GFP-RHD2 expressed under the *RHD2* promoter. However, the spatiotemporal mechanism of AtRBOHC/RHD2 subcellular redistribution and delivery to the plasma membrane (PM) during root hair initiation and tip growth are still unclear. Here, we used advanced microscopy for detailed qualitative and quantitative analysis of vesicular compartments containing GFP-RHD2 and characterization of their movements in developing bulges and growing root hairs. These compartments, identified by an independent marker such as the trans-Golgi network (TGN), deliver GFP-RHD2 to the apical PM domain, the extent of which correlates with the stage of root hair formation. Movements of TGN/early endosomes, but not late endosomes, were affected in the bulging domains of the *rhd2-1* mutant. Finally, we reveal that accumulation in the growing tip, docking, and incorporation of TGN compartments containing GFP-RHD2 to the apical PM of root hairs requires structural sterols. These results help clarify the mechanism of polarized AtRBOHC/RHD2 targeting, maintenance, and recycling at the apical PM domain, coordinated with different developmental stages of root hair initiation and growth.

**One-sentence summary:** Advanced microscopy and quantitative analysis of vesicular TGN compartments revealed that delivering GFP-RHD2 to the apical plasma membrane domains of developing bulges and growing root hairs requires structural sterols.

## Introduction

Root hairs are specialized lateral tubular extensions of root epidermal cells called trichoblasts. Although their presence is dispensable for plant survival, they considerably increase root surface area. Root hairs are involved in effective water and nutrient uptake, which is important for sustained plant growth. Root hairs represent an excellent and easily accessible model of polar cell expansion in plants because of their position at the root surface, simple structure, and fast growth. Root hair development starts with bulging at the outer tangential cell wall/plasma membrane (PM) domain in the rootward pole of the trichoblasts, followed by subsequent transition to tip growth, which is polar expansion at the tip. Tip growth ensures the formation of long tubular root hair (Carol and Dolan, 2002; Grierson and Schiefelbein, 2002).

Root hair tip growth is caused by the establishment and maintenance of cellular polarity since it requires polarization of the cytoskeleton, membrane trafficking, and localized cell wall deposition (Šamaj et al., 2006). A tip-focused gradient of Ca^2+^ ions also plays a significant role in all tip-growing cells including root hairs. Hyperpolarization-activated cation channels maintain an increase in cytosolic Ca^2+^ concentration in the growing tip, transporting apoplastic Ca^2+^ ions inwardly across the PM (Kiegle et al., 2000; Véry and Davies, 2000; Miedema et al., 2001). This tip-focused cytosolic Ca^2+^ gradient is established upon root hair tip growth initiation and persists until its cessation (Dolan et al., 1994; Wymer et al., 1997; Foreman et al., 2003). Other components of the tip growth mechanism are reactive oxygen species (ROS), recognized by calcium channels and gate their opening. These ROS are generated by the NADPH oxidase RESPIRATORY BURST OXIDASE HOMOLOG PROTEIN C (RBOHC), which is encoded by the *RBOHC/RHD2* (*ROOT HAIR DEFECTIVE 2*) locus in *Arabidopsis thaliana* (Foreman et al., 2003). The genome of *A. thaliana* encodes 10 different RBOH members (RBOHA - RBOHJ; Torres et al., 2002; Sagi and Fluhr, 2006). Nevertheless, among all RBOH family members, only AtRBOHC/RHD2 is required to regulate the early stages of root hair tip growth. Mutations in *RBOHH* and *RBOHJ* lead to defects in root hair elongation (Mangano et al., 2017). Although the root hairs of these mutants were shorter than those of the wild type, they were considerably longer in comparison to loss-of-function mutants in *RBOHC*. Moreover, the most considerable defects in *rbohh* and *rbohj* mutants are caused by disturbed pollen tube elongation, leading to reduced mutant fertility (Kaya et al., 2019; Zhou et al., 2020).

It has been shown that *RHD2* transcription is activated in the root epidermis of diverse root zones from the proximal meristem to the elongation zone (Foreman et al., 2003). Loss-of-function mutations in the *AtRBOHC/RHD2* locus in *rhd2* mutants resulted in short root hairs with missing tip-focused ROS and Ca^2+^ gradients (Schiefelbein and Somerville, 1990; Foreman et al., 2003). Importantly, root hair tip growth of *rhd2-1* mutant can be partly restored by external applications of ROS or by pH alkalization that reinstates the PM activity of Ca^2+^ channels (Monshausen et al., 2007). This indicates positive feedback between apical localization and activity of RHD2, ROS, and Ca^2+^ gradients, controlling root hair development (Foreman et al, 2003; Carol et al., 2005). A fully complemented phenotype of *rhd2* mutants with rescued root hair development has been achieved after transformation with GFP-tagged functional AtRBOHC (GFP-RHD2) under the control of its native promoter (Takeda et al., 2008). Tissue- and cell-specific localization studies revealed that GFP-RHD2 signals, particularly in trichoblasts, accumulate in the bulges and apices of growing root hairs (Takeda et al., 2008).

Root hair tip growth and development are supported by dynamic vesicular trafficking. The apical zone of growing root hairs is enriched with secretory and endocytic/recycling vesicles, balancing macromolecule supply, retrieval, and recycling (Cole and Fowler, 2006; Šamaj et al., 2006; Campanoni and Blatt, 2007). Endosomal compartments represent endomembrane trafficking hubs categorized into early endosomes/*trans*-Golgi networks (TGN) and late endosomes/multivesicular bodies (Reyes et al., 2011). Furthermore, early endosomes/TGN compartments merge secretory and endocytic pathways (Viotti et al., 2010; Contento and Bassham, 2012; Qi and Zheng, 2013), and late endosomes/multivesicular bodies are involved in transport pathways towards vacuoles (Bottanelli et al., 2011; Contento and Bassham, 2012). Rab GTPases in the clear zone of root hairs spatiotemporally organize vesicular trafficking, exocytosis, endocytosis and membrane recycling (Voigt et al., 2005; Berson et al., 2014).

FM dyes allow the fluorescence labeling of PM, highly dynamic populations of early, recycling, and late endosomes in root hairs (Ovečka et al., 2005). Detailed analysis and characterization of distinct endosomal compartments have been achieved using protein markers translationally fused to fluorescent proteins; early endosomes/TGN compartments were specifically detected by RabA1d and VTI12, a SNARE-protein (Sanderfoot et al., 2001; Uemura et al., 2004; Ovečka et al., 2010; Berson et al., 2014). Molecular marker based on the FYVE domain, specifically binding to phosphoinositol-3-phosphate (PI-3P), was used to detect late endosomes (Gillooly et al., 2001; Voigt et al., 2005). Members of the Rab-GTPase family have been verified as markers for late endosomal compartments and eventually colocalize with FYVE, such as RabF2a, RabF1 (Ueda et al., 2004; Voigt et al., 2005; Haas et al., 2007), and RabF2b (Geldner et al., 2009). The dynamic properties of early and late endosomes using molecular markers have been described in the growing root hairs of Arabidopsis (von Wangenheim et al., 2016).

Structural sterols modulate the permeability and fluidity of plant membranes and influence the physical and physiological properties of membrane proteins (Mukherjee and Maxfield, 2000; Clouse, 2002). They play important roles in vesicle trafficking and docking, signaling, and protein localization in membranes (Lindsey et al., 2003), particularly in tip-growing root hairs (Ovečka et al., 2010) and pollen tubes (Liu et al., 2009). Sterols interact with phospholipids and create macromolecular nanodomains in the extracellular leaflet of the PM, so-called “lipid rafts” (Simons and Toomre, 2000). Lipid rafts are involved in the perception and transduction of signals associated with membrane receptors, selecting exo-endocytic cargo molecules, and intracellular redistribution of sterols (Menon, 2002). Proteomic studies identified NADPH oxidases NtRBOHD, StRBOHB and AtRBOHB with other PM proteins enriched in detergent-resistant membrane fraction of suspension cells (Morel et al., 2006, Srivastava et al., 2013). Subcellular visualization revealed AtRBOHD localization to dynamic spots at the PM and its endocytosis induced by salt stress (Hao et al., 2014). Nevertheless, the spatiotemporal relationship between local arrangements of structural sterols and integral PM proteins such as AtRBOHC/RHD2 during root hair development is still unclear.

This study examined the qualitative and quantitative localization patterns of GFP-RHD2 during root hair formation in Arabidopsis using advanced microscopy methods. We revealed that incorporating GFP-RHD2 into the apical and subapical PM domains was tightly correlated with the establishment of the apical zone in bulges and growing root hairs. We documented the delivery of GFP-RHD2 to the apical zone during bulge formation and the apex of growing root hairs by dynamic vesicular compartments. These were identified as early endosomal/TGN compartments using colocalization analyses with reliable endosomal molecular markers. Single-particle tracking analysis revealed complex movement patterns of vesicular compartments containing GFP-RHD2 and identified their interactions with the apical PM during root hair formation. In addition, we revealed the role of structural sterols in the docking of GFP-RHD2 vesicles at the apical PM domain of root hairs. Consequently, the genetic approach showed that early endosomal/TGN compartments were more seriously affected in their movements than late endosomes in the *rhd2-1* mutant. This quantitative advanced microscopy study showed the complex nature of GFP-RHD2 delivery, maintenance, and recycling at the apical PM, characteristic of root hair formation in Arabidopsis.

## Results

### Cell type-specific and subcellular distribution pattern of GFP-RHD2 in Arabidopsis root

Light-sheet fluorescence microscopy (LSFM) provided visualization for developmentally regulated and cell type-specific distribution patterns of GFP-RHD2 in growing Arabidopsis roots. The GFP-RHD2 signal was observed in epidermal cells in root hair formation zone, and was explicitly accumulated in the bulges and apices of developing root hairs (Figure 1A). Orthogonal projections from different root developmental zones confirmed GFP-RHD2 localization in the root epidermis, preferably in trichoblasts within the root differentiation and elongation zone (Figure 1B). Specifically, GFP-RHD2 accumulated at the tips of emerging (Figure 1A and C) and growing (Figure 1A and D) root hairs. This distribution and accumulation pattern of GFP-RHD2 was confirmed by a pseudo-color-coded, semi-quantitative fluorescence intensity evaluation for maximum intensity projections of the whole root apex (Figure 1A; Supplemental Video S1) and orthogonal projections of different root developmental zones (Figure 1B-D). Advanced cellular and sub-cellular localization of GFP-RHD2 using Airyscan CLSM revealed differential abundance of GFP-RHD2 in trichoblasts and atrichoblasts (Figure 1E) as well as specific enrichment in bulge selection sites and developing bulges of trichoblasts (Figure 1E and F). Most importantly, Airyscan CLSM imaging revealed GFP-RHD2 presence in small motile compartments in the cytoplasm (Figure 1E and F). Their number was considerably higher in trichoblasts (Figure 1E), and there was an apparent progressive accumulation in developing bulges (arrowhead in Figure 1E and F). In addition to the visualization of GFP-RHD2 in small compartments in the cytoplasm, Airyscan CLSM revealed localization in the PM, particularly at bulging sites and apical domains of developing bulges (Figure 1E and F).

**Figure 1.**
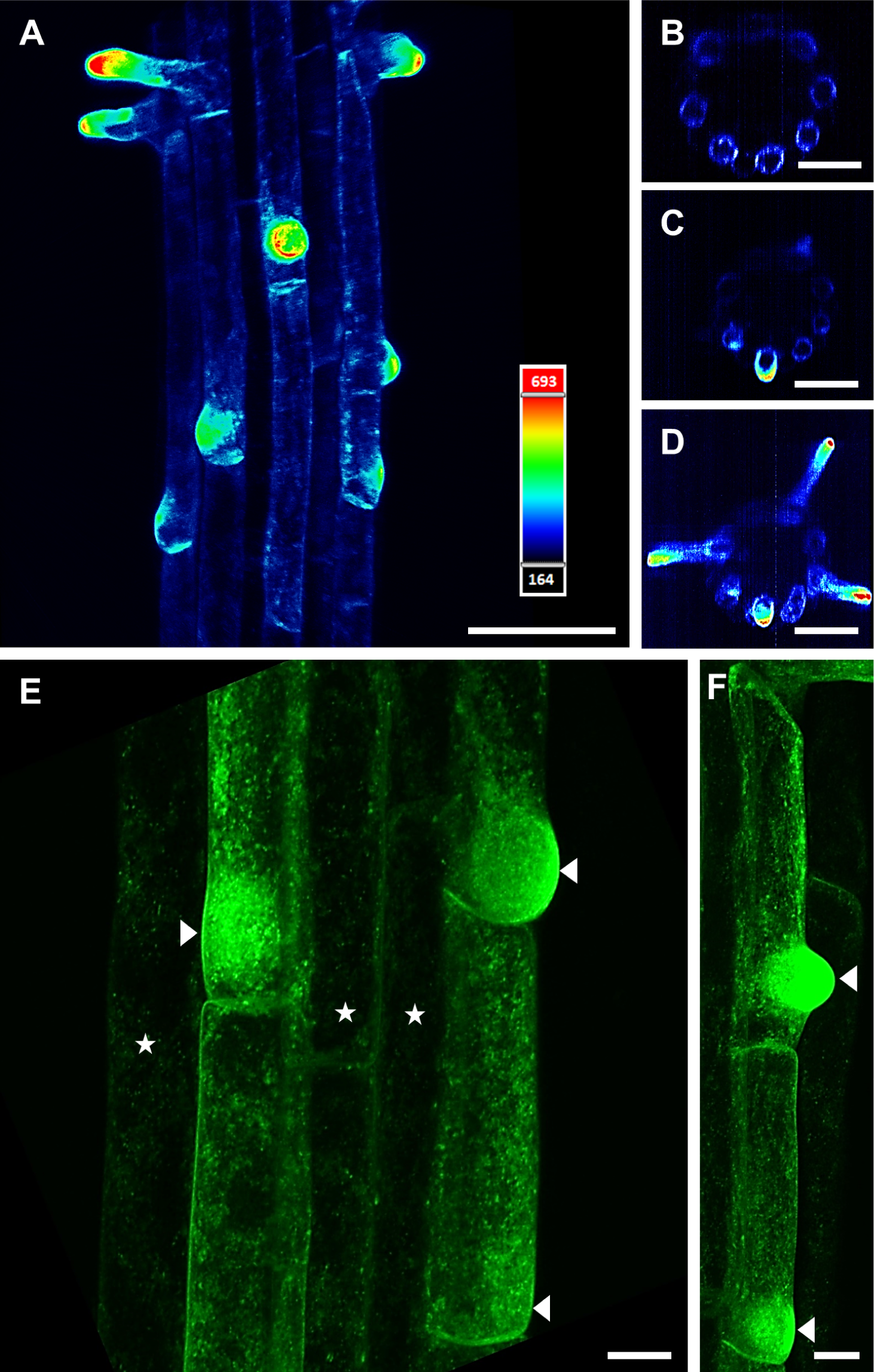
Expression pattern and cell type-specific distribution of GFP-RHD2 in growing Arabidopsis root. **(A-D)** Distribution pattern in root hair formation zone **(A)**, where GFP-RHD2 was expressed mainly in trichoblasts **(B)** and accumulated at tips of emerging **(C)** and growing **(D)** root hairs. Maximum intensity projection of the root hair formation zone **(A)** and orthogonal projections at different developmental root zones **(B-D)** from light-sheet fluorescence microscopy imaging as visualized using semi-quantitative fluorescence intensity distribution and pseudo-color-coded scale, where dark blue represents minimal intensity (164 arbitrary units) and pale red represents maximum intensity (693 arbitrary units). Spatial overview of the maximum intensity projection from the root hair formation zone **(A)** is presented in **Supplemental Video S1**. **(E, F)** Detailed visualization of subcellular compartments containing GFP-RHD2 in trichoblasts and their progressive accumulation in developing bulges (arrowheads) using Airyscan confocal laser scanning microscopy. A substantially lower number of compartments with GFP-RHD2 was detected in atrichoblasts (asterisks). Scale bar = 50 µm **(A)**, 40 µm **(B-D)**, 10 µm **(E, F)**.

We used time-lapse imaging of dynamic GFP-RHD2 localization and targeted accumulation during root hair formation to present the whole dynamic process in the form of a video. The localization pattern of GFP-RHD2 can be related to the root hair growth rate, which is known to change in diverse root hair developmental stages dynamically. Time-lapse recording (Supplemental Video S2) allowed identification of particular stages from early bulges (stage I; Figure 2A) to late bulges (stage II; Figure 2B) at the beginning, followed by subsequent stages of accelerated root hair tip growth. These subsequent stages were represented by short root hairs (stage III; Figure 2C), a transition from short to long root hairs (stages III-IV; Figure 2D), and long root hairs (stage IV; Figure 2E). The pattern of the tip growth rate over the entire process was quantitatively analyzed using kymographs. In the long-term time-lapse acquisition set up, which was based on recording every 5 min (Figure 2A-E; Supplemental Video S2), we recorded a slow extension in early bulges (stage I) and a faster extension in late bulges (stage II; Supplemental Figure S1A and B), followed by fast, smooth, and continuous tip growth of short and long root hairs (stages III and IV; Supplemental Figure S1C-F). Non-growing root hairs (Supplemental Figure S1G) were also identified using this method (Supplemental Figure S1H). The slow mode of the bulge extension versus the fast mode of the root hair extension and particularly sustained regular mode of tip growth in short and long root hair stages were revealed (Figure 2F). Sub-cellular time-lapse imaging of GFP-RHD2 distribution at different stages of root hair initiation showed polarized accumulation at the bulging sites and emerging bulges. GFP-RHD2 positive compartments were particularly enriched in the cortical cytoplasm of the bulging domain and selectively targeted the apical PM of the emerging bulge (Figure 2G). Polarized accumulation of GFP-RHD2 at the apical PM was documented in the late bulges (Figure 2H and I; Supplemental Video S3), short (Figure 2J and K; Supplemental Video S4), and long (Figure 2L and M; Supplemental Video S5) growing root hairs. GFP-RHD2 at the apical PM was temporarily maintained also in growth-terminating root hairs (Figure 2N and O; Supplemental Video S6). Importantly, GFP-RHD2 was present in the cortical cytoplasm of developing bulges (Figure 2H and I; Supplemental Video S3) and accumulated in the apical, vesicle-rich “clear” zone of tip-growing root hairs (Figure 2J-M; Supplemental Videos S4 and S5). Thus, GFP-RHD2 accumulation in PM was detected only in specific domains of the trichoblast, bulging out of the parental outer tangential PM during bulge formation (Figure 2A and B) and in the apex of growing root hairs (Figure 2C-E; Supplemental Video S2). Live cell imaging revealed the presence of GFP-RHD2 in small compartments of the cortical cytoplasm (Figure 2G) and targeted GFP-RHD2 accumulation in the apical PM. Visualization of GFP-RHD2 distribution in growing root hairs, both at early (Figure 2J and K) and later (Figure 2L and M) stages, showed an accumulation of compartments containing GFP-RHD2 in the apical PM, clear zone, and subapical cytoplasm. Similarly, the PM specifically accumulated GFP-RHD2 only within the apical and subapical zones of root hairs (Figure 2J-M; Supplemental Figure S1C and E).

**Figure 2.**
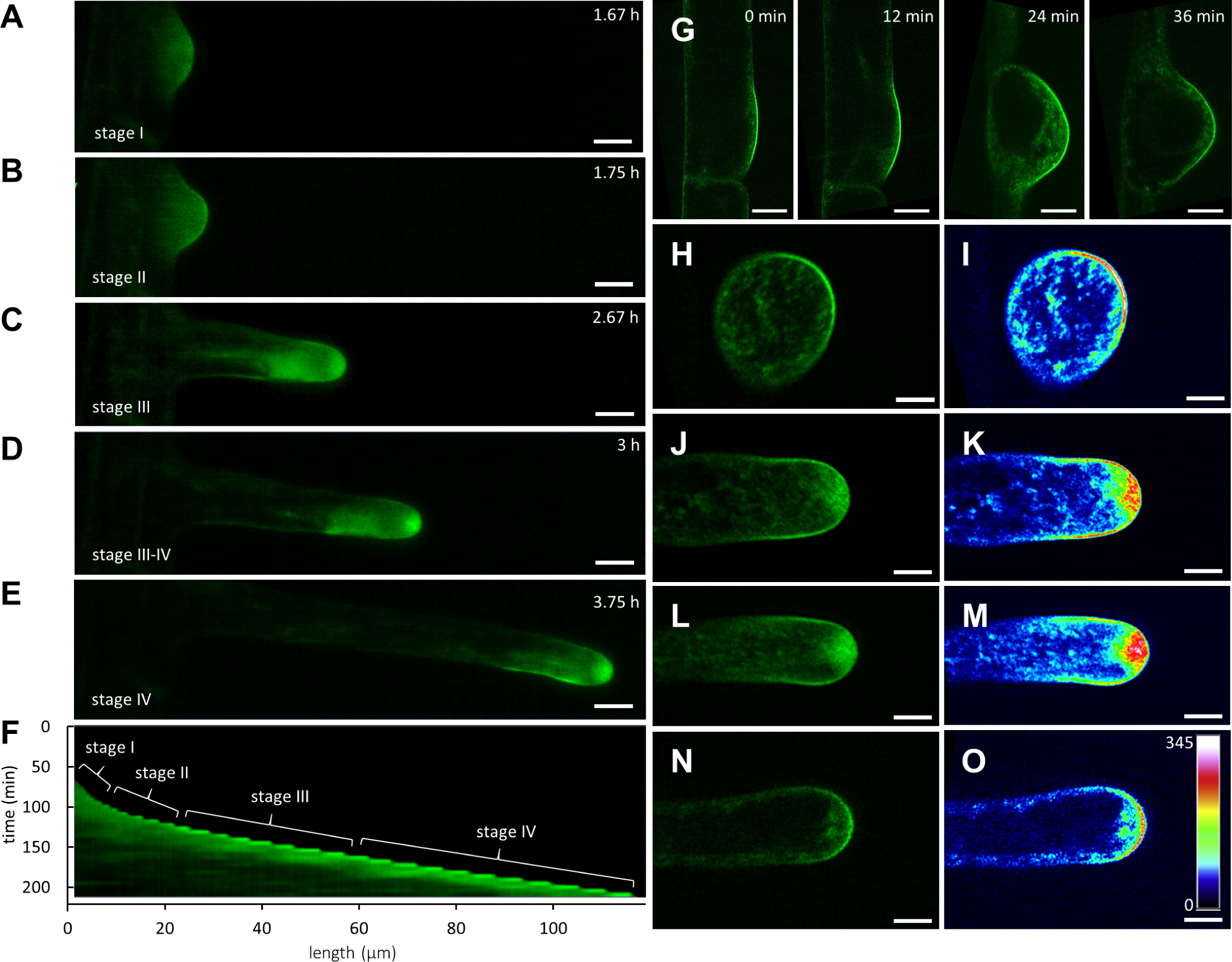
Time-course imaging of GFP-RHD2 localization and accumulation during root hair formation. **(A-F)** Localization pattern of GFP-RHD2 in different stages of root hair development and its relation to particular root growth rate. Time-lapse recording of developing root hair showing stages of early bulge (**A**; stage I), late bulge (**B**; stage II), short root hair (**C**; stage III), transition from short to long root hair (**D**; stages III-IV) and long root hair (**E**; stage IV). The whole process shown in **Supplemental Video S2**, is quantitatively presented in kymograph **(F)**, representing the pattern and rate of the growth of this particular root hair, growing over the distance of 120 µm within the time period of 220 min. Individual developmental stages presented in **(A-E)** are depicted in **(F)**. **(G)** Lateral view on the polarized accumulation of GFP-RHD2 at the plasma membrane domain and cortical cytoplasm of the bulging site and during the emerging of the bulge. **(H-O)** The GFP-RHD2 distribution **(H, J, L, N)** and a semi-quantitative signal intensity visualization using a pseudo-color-coded range **(I, K, M, O)** of the polarized accumulation at the apical plasma membrane domain and cortical cytoplasm in the late bulge **(H, I, Supplemental Video S3)**, short **(J, K, Supplemental Video S4)** and longer **(L, M, Supplemental Video S5)** growing root hairs, and in non-growing root hair **(N, O, Supplemental Video S6)**. Pseudo-color-coded scale shows a range from dark blue representing minimal intensity (0 arbitrary units) to white representing maximum intensity (345 arbitrary units). Timing of the sequential imaging is indicated in hours (h) using light-sheet microscopy **(A-E)** and in minutes (min) using Airyscan confocal laser scanning microscopy **(G)**. Scale bar = 10 µm **(A-E)**, 5 µm **(G-O)**.

Localization analysis showed a different range of GFP-RHD2 specific targeting to the apical PM domain at distinct stages of root hair development. Therefore, we analyzed the range of PM decorated with GFP-RHD2 (Figure 3A). A quantitative evaluation revealed that the extent of the PM zone containing GFP-RHD2 increased progressively from early to late bulges. This zone was enlarged in short-growing root hairs and reached a maximum in longer-growing root hairs but declined considerably in non-growing root hairs (Figure 3B). Thus, the distribution pattern of GFP-RHD2 at the apical PM was tightly correlated with the individual developmental stages of root hairs (Figure 3A and B). This finding was further corroborated by the GFP-RHD2 fluorescence signal intensity distribution. Visualization using the semi-quantitative 2.5-D rendering function demonstrated a gradual increase in GFP-RHD2 fluorescence signal intensity in successive root hair developmental stages, from the development of the early bulge up to fast-growing root hairs (Supplemental Figure S2).

**Figure 3.**
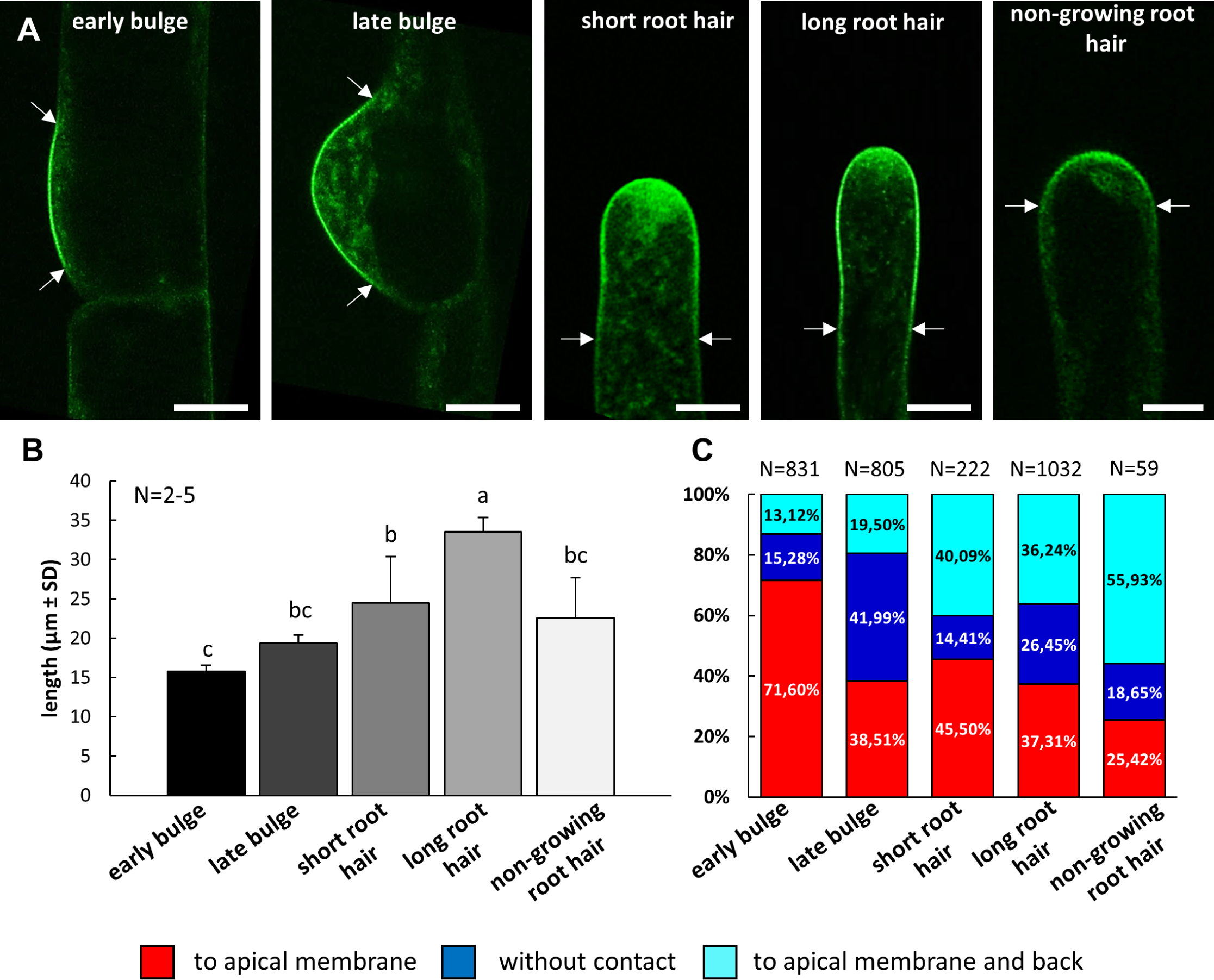
Determination of GFP-RHD2 localization at the apical plasma membrane domain at different stages of root hair formation and growth. **(A)** Delineation of the area (by two arrows) where GFP-RHD2 was specifically targeted to the apical plasma membrane domain at early and late bulging stage, in short and longer growing root hairs, and in non-growing root hairs. **(B)** Quantitative evaluation of the apical plasma membrane domain length decorated by GFP-RHD2 at different stages of root hair development. Measurement was done from 2-5 root hairs. Different lowercase letters indicate statistical significance between treatments (P < 0.05). **(C)** Quantitative determination of the ratio between the compartments containing GFP-RHD2 and showing targeted movement to the apical plasma membrane (red), moving in the cytoplasm without any contact with the apical plasma membrane (blue), or moving to and contacting the plasma membrane and subsequently moving back to the cytoplasm (cyan). The GFP-RHD2 positive compartments were tracked and analyzed at denoted stages of root hair development as depicted in **(A)**. More than 59 compartments from 3 to 7 growing root hairs from individual plants were analyzed. Scale bar = 5 µm **(A)**.

Live cell imaging revealed targeted delivery of GFP-RHD2 in dynamic vesicular compartments to the cortical cytoplasm and apical PM of developing root hairs. Based on the nature of their movements, we determined three groups of vesicular compartments carrying GFP-RHD2. The first group was represented by compartments with targeted movement to the apical PM. The second group consisted of compartments moving in the cytoplasm without any contact with the apical PM. The third group of compartments moved towards the PM and subsequently moved back to the cytoplasm after making contact. The quantitative evaluation showed a high proportion of GFP-RHD2 vesicular compartments moving towards the apical PM and a low number of compartments moving without contact or moving back after contact with the PM in the early bulge stage (Figure 3C). However, this turned to different pattern during root hair development. The compartments from the first group were relatively high in short and longer growing root hairs, while the compartments from the third group considerably increased (Figure 3C). With the cessation of tip growth, there was a considerable reduction in compartments moving to the apical PM (Figure 3C). This analysis showed that the extent of GFP-RHD2 specifically targeted to the apical PM correlated with the stages of root hair formation. Importantly, this correlation was directly affected by the ratio between GFP-RHD2 vesicular compartments targeted to the apical PM and those moving back to the cytoplasm after contacting the PM.

### Identification of GFP-RHD2 positive compartments

Next, we analyzed the nature of compartments containing GFP-RHD2 by colocalization with the selective membrane styryl dye FM4-64. In root hairs, the FM4-64 dye is able to stain the PM and early endocytotic/recycling vesicular compartments after a minute-long exposure period (Ovečka et al., 2005). After 10 min of application, FM4-64 colocalized with GFP-RHD2 at the PM and in motile vesicular compartments in the apical and subapical zones of growing root hairs (Figure 4A). The colocalization at PM was semi-quantitatively documented by a cross-sectional fluorescence intensity profile measurement within the subapical zone of the root hair (Figure 4B). A high degree of colocalization in moving vesicular compartments within the subapical zone of growing root hairs (Figure 4C) was revealed by fluorescence intensity profile measurements (Figure 4D). Semi-quantitative measurement of fluorescence signal intensities of GFP-RHD2 and FM4-64 at the PM along the whole longitudinal root hair axis showed a stable level of FM4-64 fluorescence in the subapical and shank regions, but GFP-RHD2 fluorescence was specifically enriched only in the apical and subapical PM domains and gradually decreased in the shank region (Figure 4E). Results from semi-quantitative profile measurements were corroborated by the quantitative colocalization analysis showing a high degree of Pearson’s correlation coefficients between FM4-64 and GFP-RHD2 in intracellular compartments (Figure 4F; Supplemental Figure S3A and B) and in the subapical PM (Figure 4G; Supplemental Figure S3C and D). This analysis suggests that vesicular compartments containing GFP-RHD2 might belong to the early endosome group.

**Figure 4.**
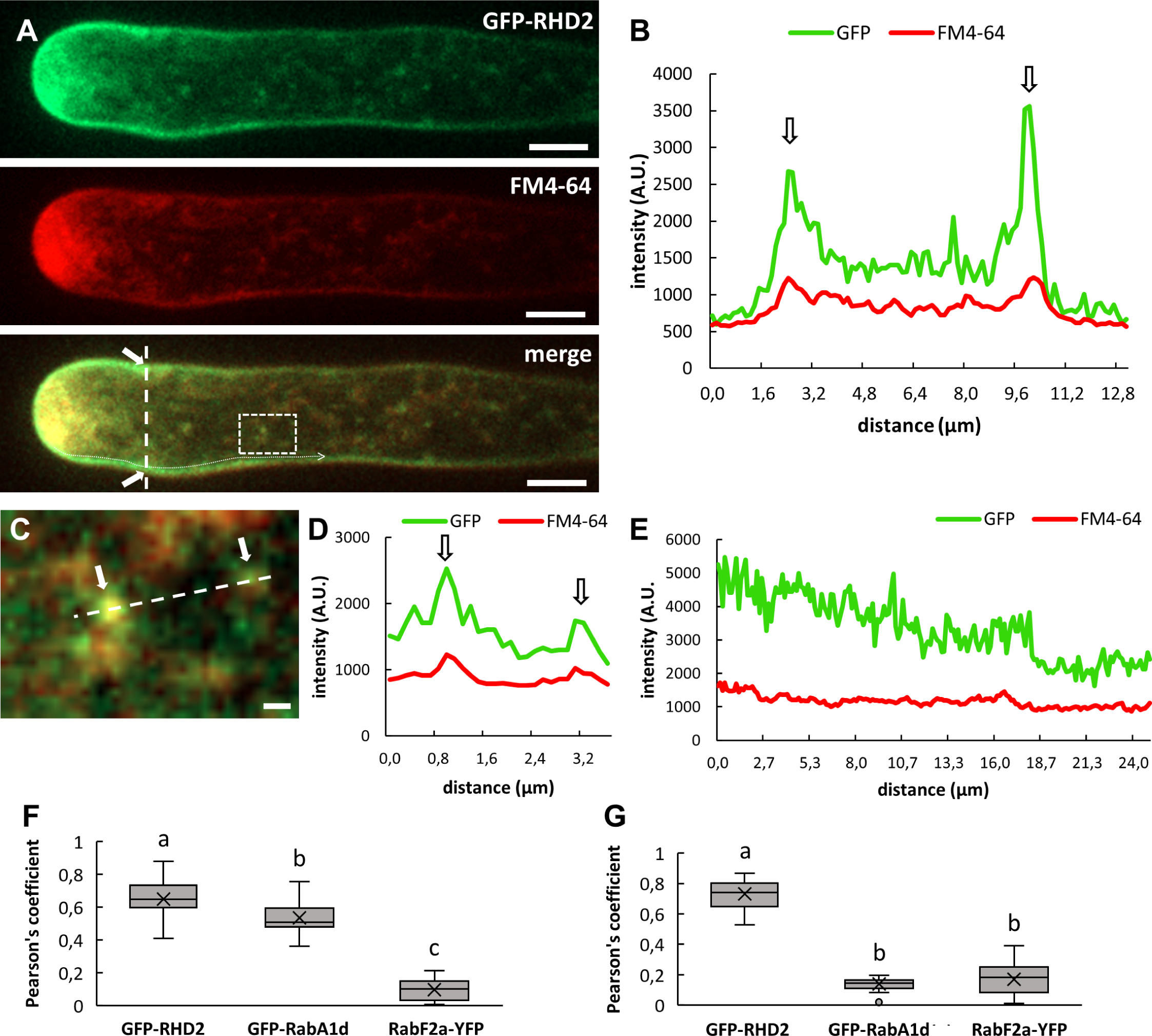
Colocalization analysis of GFP-RHD2 with selective membrane styryl dye FM4-64 in growing root hairs. **(A)** Localization pattern of GFP-RHD2 in the apical plasma membrane domain, cytoplasm of the apical and subapical zone, and in motile compartments of growing root hair co-labeled with membrane styryl dye FM4-64. Colocalization analysis has been performed in merged image along the cross-section profile marked by interrupted white line (white arrows indicate the plasma membrane), in compartments within the white box, and along white dotted line. **(B)** Fluorescence intensity profile of the GFP and FM4-64 signal measured along the cross-section interrupted white line in **(A)**. Arrows indicate position of the plasma membrane. **(C)** Detailed image of compartments containing GFP-RHD2 and FM4-64 within the white box depicted in **(A)**. Arrows indicate position of the compartments and interrupted white line depicts the profile for fluorescence intensity quantification. **(D)** Fluorescence intensity profile of the GFP and FM4-64 signal along the interrupted white line in **(C)**. Arrows indicate position of the compartments. **(E)** Fluctuation of the GFP and FM4-64 fluorescence signal intensity at the plasma membrane along the longitudinal root hair axis from apex to the shank region measured along the white dotted line in **(A)**. Note the stable level of FM4-64 fluorescence signal intensity at the plasma membrane in subapical and shank region, while GFP-RHD2 fluorescence signal intensity was specifically enriched in the apical and subapical plasma membrane domains and gradually decreased within the shank region. **(F, G)** Quantitative colocalization analysis using Pearson’s correlation coefficients between FM4-64 and GFP-RHD2, early endosomal marker GFP-RabA1d **(Supplemental Figure S4A)**, and late endosomal marker RabF2a-YFP **(Supplemental Figure S4B)** in intracellular compartments **(F)**, and in subapical plasma membrane **(G)**. Selections of ROIs and scatterplots of measurements are presented in **Supplemental Figures S3, S5, S6**. Scale bar = 5 µm **(A)**, 1 µm **(C)**.

Transgenic Arabidopsis line carrying GFP-RabA1d marker was used to visualize early endosomes/TGN compartments (Ovečka et al., 2010; Berson et al., 2014; von Wangenheim et al., 2016). Colocalization analysis of GFP-RabA1d with FM4-64 in growing root hairs revealed no overlap at the PM (Supplemental Figure S4A and C). However, GFP-RabA1d colocalized with FM4-64 (applied for 10 min) in motile compartments (Supplemental Figure S4E and F). This was corroborated by a quantitative colocalization analysis using Pearson’s correlation coefficients, showing high values in intracellular compartments (Figure 4F; Supplemental Figure S5A and B) and low values in the subapical PM (Figure 4G; Supplemental Figure S5C and D). RabF2a-YFP is a good marker of late endosomes (von Wangenheim et al., 2016) and colocalization analysis with FM4-64 (applied for 10 min) revealed no colocalization in growing root hairs both at the PM (Supplemental Figure S4B and D) and in motile compartments (Supplemental Figure S4G and H). Again, a quantitative colocalization analysis using Pearson’s correlation coefficients corroborated this observation, showing very low values both in intracellular compartments (Figure 4F; Supplemental Figure S6A and B) and in the subapical PM (Figure 4G; Supplemental Figure S6C and D).

Colocalization analysis using FM4-64 suggested that vesicular compartments delivering GFP-RHD2 towards the apical PM in growing root hairs are likely early endosomes/TGN. To confirm this suggestion, we prepared double transgenic lines carrying GFP-RHD2 together with the early endosomal/TGN marker, mCherry-VTI12, or with the late endosomal marker, mCherry-RabF2b. Markers mCherry-VTI12 and mCherry-RabF2b were not detected in the PM of growing root hairs (Supplemental Figure S7). Therefore, these two markers did not colocalize with GFP-RHD2 at the PM (Figure 5A-D and J; Supplemental Figures S8C and D; S9C and D). However, the early endosomal/TGN marker mCherry-VTI12 showed colocalization with GFP-RHD2 in motile compartments (Figure 5A and E); this was confirmed by fluorescence intensity profile quantification (Figure 5F) and a quantitative colocalization analysis using Pearson’s correlation coefficients (Figure 5I; Supplemental Figure S8A and B). Topological colocalization analysis (Figure 5B), fluorescence intensity profile quantification (Figure 5G and H), and quantitative colocalization analysis using Pearson’s correlation coefficients (Figure 5I; Supplemental Figure S9A and B) suggested no colocalization between GFP-RHD2 and mCherry-RabF2b, which represents marker for late endosomal compartments. Concluding from colocalization results with FM4-64 and endosomal markers, vesicular trafficking and targeted delivery of GFP-RHD2 during root hair development depend on early endosomal/TGN compartments.

**Figure 5.**
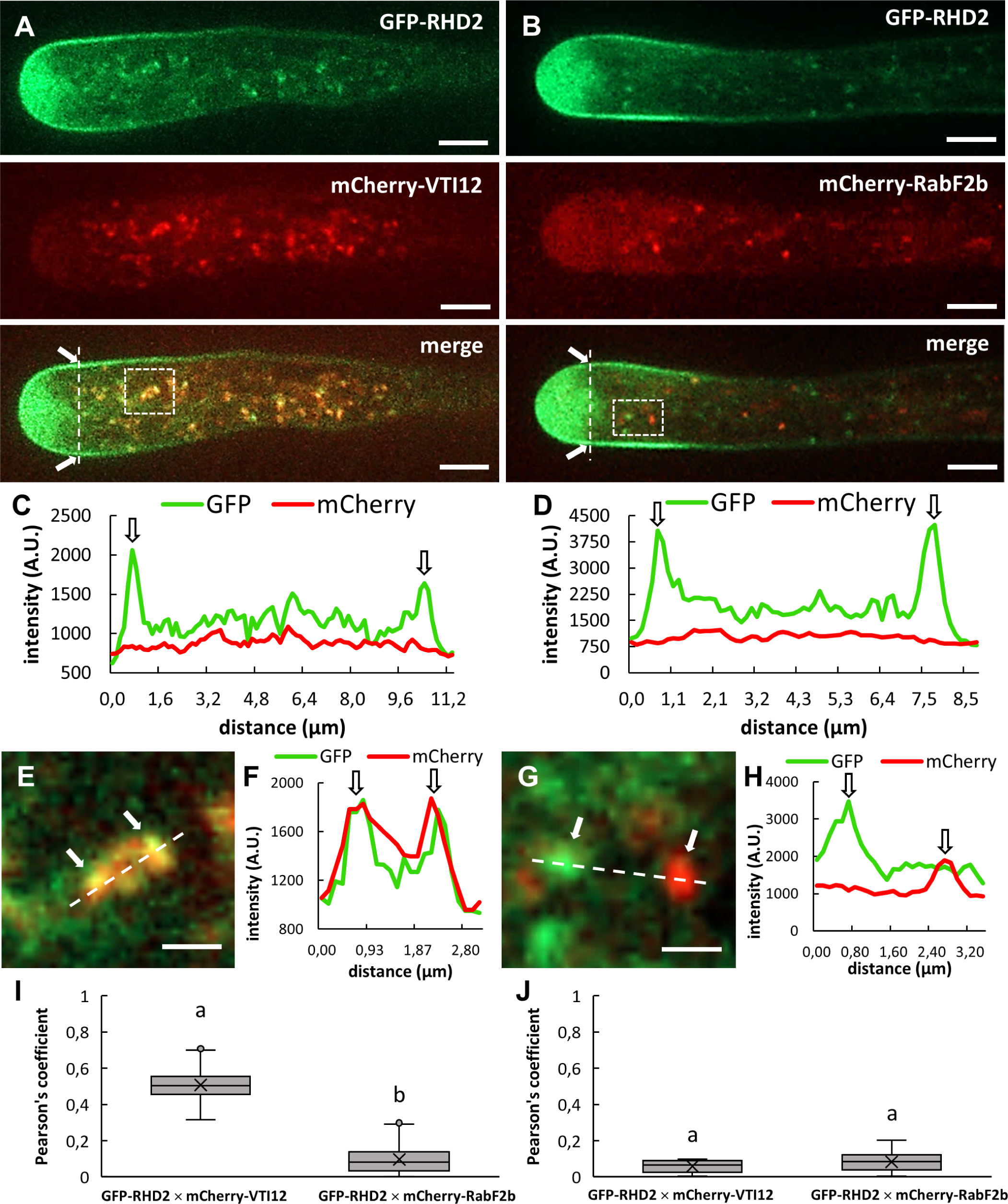
Colocalization analysis of GFP-RHD2 with early and late endosomal markers in growing root hairs. **(A)** Distribution of GFP-RHD2 in the apical plasma membrane domain, the clear zone, cytoplasm of the subapical zone, and in moving compartments of growing root hair of Arabidopsis double transgenic line carrying both GFP-RHD2 and the early endosomal/trans-Golgi network marker mCherry-VTI12. **(B)** Distribution of GFP-RHD2 in the apical plasma membrane domain, the clear zone, cytoplasm of the subapical zone, and in moving compartments of growing root hair of Arabidopsis double transgenic line bearing both GFP-RHD2 and the late endosomal marker mCherry-RabF2b. In merged images of **A** and **B**, an extent of fluorescence intensity colocalization has been analyzed along the cross-section profiles marked by interrupted white lines (white arrows indicate the plasma membrane) and in moving compartments within the white boxes. **(C)** Fluorescence intensity profiles of GFP and mCherry signals along the cross-section interrupted white line of the root hair shown in **(A)**. **(D)** Fluorescence intensity profiles of the GFP and mCherry signal along the cross-section interrupted white line of the root hair shown in **(B)**. Arrows in **C** and **D** indicate position of the plasma membrane. **(E)** Detailed image of compartments containing GFP-RHD2 and mCherry-VTI12 within the white box in **(A)**. **(F)** Fluorescence intensity profile of the GFP and mCherry-VTI12 signal along the interrupted white line in **(E)**. **(G)** Detailed image of compartments positive to GFP-RHD2 and mCherry-RabF2b within the white box in **(B)**. **(H)** Fluorescence intensity profile of the GFP and mCherry-RabF2b signal along the interrupted white line in **(G)**. Arrows indicate position of the compartments. **(I, J)** Quantitative colocalization analysis using Pearson’s correlation coefficients between GFP-RHD2 and the early endosomal/trans-Golgi network marker mCherry-VTI12, and between GFP-RHD2 and the late endosomal marker mCherry-RabF2b, in intracellular compartments **(I)**, and in subapical plasma membrane **(J)**. Selections of ROIs and scatterplots of measurements are presented in **Supplemental Figures S8-S9**. Scale bar = 5 µm **(A, B)**, 1 µm **(E, G)**.

### Quantitative microscopic analysis of motile compartments containing GFP-RHD2 at different stages of root hair development

Live cell imaging of developing root hairs by advanced microscopy enabled the qualitative determination of compartments containing GFP-RHD2. We analyzed the quantitative parameters of their movements in different stages of root hair formation using the particle-tracking method and characterized three independent groups of the GFP-RHD2 compartments. The first group was labeled in red (compartments moving to the apical PM and fusing with it), the second group was labeled in blue (compartments moving in the cytoplasm without any contact with the apical PM), and the third group was labeled in cyan (compartments moving to and contacting the PM and subsequently moving back to the cytoplasm). Movement distances of GFP-RHD2 compartments during a 60 s period imaged by Airyscan CLSM were tracked using the “tracked cells function” of the ARIVIS Vision4D software. These trajectories were directly displayed in a color-coded mode in the image sequences of early (Figure 6A), and late (Figure 6C) bulges, as well as short (Figure 6E) and longer (Figure 6G) growing root hairs. In addition, the values of the maximum and the average speed of movement were analyzed.

**Figure 6.**
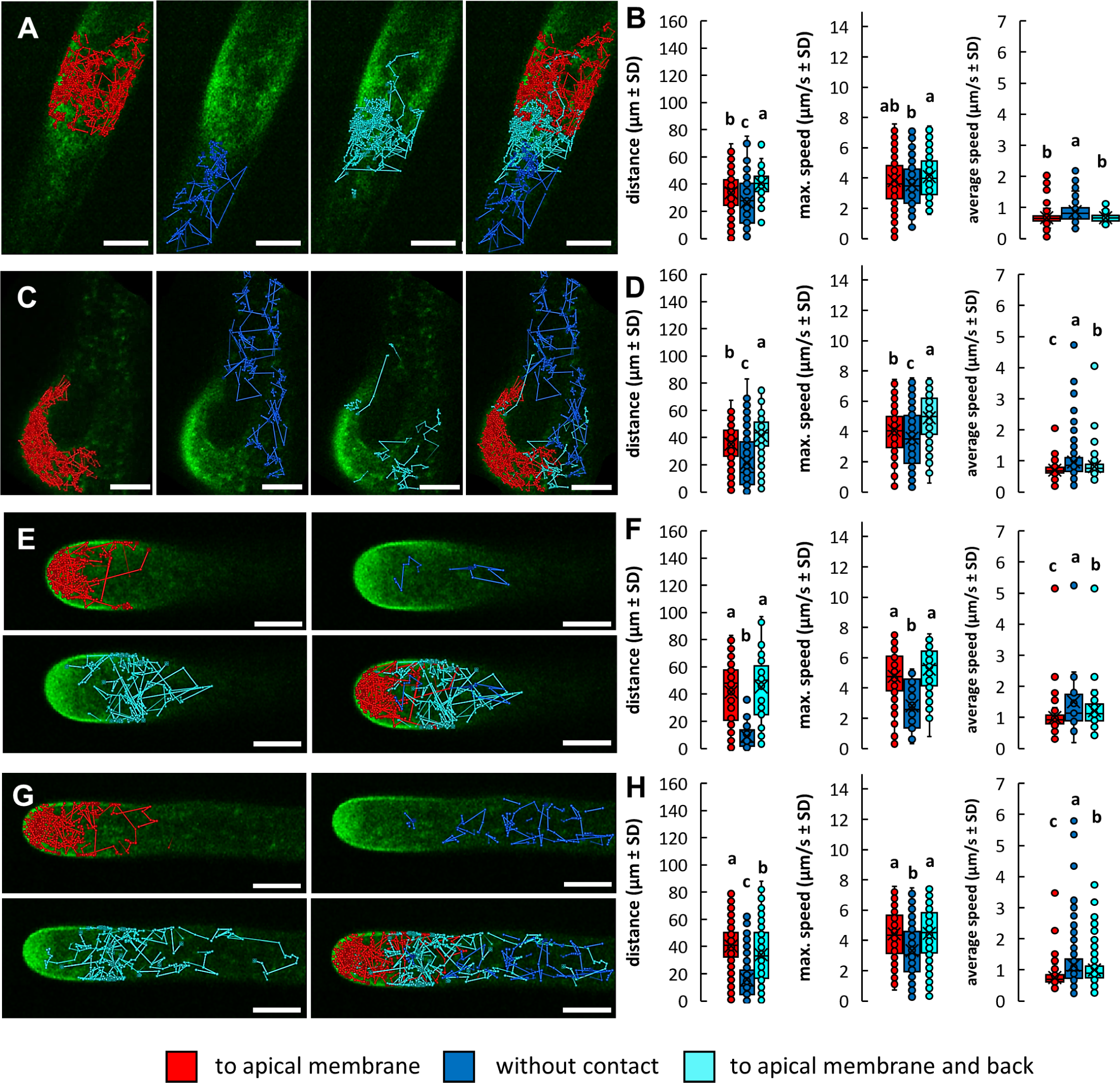
Qualitative determination and quantitative tracking of moving compartments containing GFP-RHD2 at different stages of root hair development. **(A, C, E, G)** Line tracking of the compartments showing targeted movement to the apical plasma membrane (red), moving in the cytoplasm without any contact with the apical plasma membrane (blue), and moving to and contacting the plasma membrane and moving back to the cytoplasm (cyan), and merged images of all three types of tracks in early bulge **(A)**, late bulge **(C)**, short **(E)**, and longer **(G)** growing root hairs. **(B, D, F, H)** Quantitative determination of tracked distance of compartments, maximum and average speeds of their movements during tracking analysis, as compared separately for the compartments showing targeted movement to the apical plasma membrane (red), moving in the cytoplasm without any contact with the apical plasma membrane (blue), and moving to and contacting the plasma membrane and moving back to the cytoplasm (cyan) in early bulge **(B)**, late bulge **(D)**, short **(F)**, and longer **(H)** growing root hairs. More than 222 compartments from 3 to 7 growing root hairs from individual plants were analyzed. Box plots show first and third quartiles, split by the median (line) and mean (cross). Whiskers extend to include the max/min values. Different lowercase letters indicate statistical significance between treatments (P < 0.05). Scale bar = 5 µm.

Line tracking trajectories in different stages of root hair development clearly distinguished spatial separation of GFP-RHD2 compartments from different groups (Figure 6A and C and E and G). Compartments from the red group were located almost exclusively in the bulging site (Figure 6A) and the whole bulges (Figure 6C) as well as in the clear and subapical zones of growing root hairs (Figure 6E and G). In contrast, GFP-RHD2 compartments from the cyan group were located mainly at the periphery of the bulging zone (Figure 6A) and developing bulging domains (Figure 6C). In growing root hairs, compartments from the cyan group moved preferably in the subapical and shank regions, contacting the lateral PM (Figure 6E and G) exclusively. Differences in the spatial distribution of individual compartments might be correlated with the distance and speed in different root hair developmental stages. Compartments from the blue group showed the shortest displacement distances in almost all cases (Figure 6B and D and F and H; Supplemental Figure S10), with the lowest values in maximum movement speed. However, the average speed of these compartments was the highest (with the highest variability) at all stages of root hair development (Figure 6B and D and F and H). Compartments from the red group moved along short distances in early and late bulging stages (Figure 6B and D) and these distances increased significantly during root hair tip growth in comparison to the cyan group (Figure 6F and H). Compartments from the cyan group moved over the longest distance in early and late bulging stages (Figure 6B and D), but these values decreased in the growing root hair stage (Figure 6F and H; Supplemental Figure S10). The maximum movement speed was comparable between compartments from the red and cyan groups, but the average speed was significantly higher in compartments from the cyan group, particularly in growing root hairs (Figure 6B and D and F and H). The trend of faster movement of compartments from the cyan group was evident from the frequency distribution graphs, where the curves for the maximum speed of cyan group compartments were right-shifted towards higher µm/s values (Supplemental Figure S10).

GFP-RHD2 was also localized to motile compartments in atrichoblasts. To characterize their properties, we carried out a comparative quantitative tracking analysis with compartments from trichoblasts moving outside of the bulging zone (Supplemental Figure S11A and B). In principle, there were no significant differences in the average distance and minimum/maximum/average movement speed (Supplemental Figure S11C and D). When the motions and dynamic behavior of compartments containing GFP-RHD2 were compared between tricho- and atrichoblasts (Supplemental Figure S11A and B), those in atrichoblasts showed more variable movement and dynamics. This trend was obvious from the frequency distribution graphs, revealing a slightly shifted distribution to longer distances for GFP-RHD2 compartments in trichoblasts (Supplemental Figure S11E). However, the frequency distribution of maximum (Supplemental Figure S11F) and average (Supplemental Figure S11G) movement speed were very similar. These data indicate that trichoblast compartments with GFP-RHD2, which are not involved in root hair formation, move similarly to those from the atrichoblasts but distinctly from those from developing root hairs.

Live cell imaging using Airyscan CSLM allowed the visualization of close contacts between motile compartments and the apical PM in the root hair and provided significantly improved spatial resolution. Time-course visualization of individual compartments containing GFP-RHD2 revealed direct contact with the PM enriched with GFP-RHD2 (Figure 7A; Supplemental Video S7). It was possible to document the full incorporation of these compartments into the apical PM by complete fusion (Figure 7B and D; Supplemental Video S8). Alternatively, we observed only temporal contacts of such compartments, lasting for ∼30 - 50 seconds, but without full integration to the PM (Figure 7C and E; Supplemental Video S9). Such docking events without complete incorporation into the PM lasted longer than the total fusion events (cf. Figure 7B and D to Figure 7C and E).

**Figure 7.**
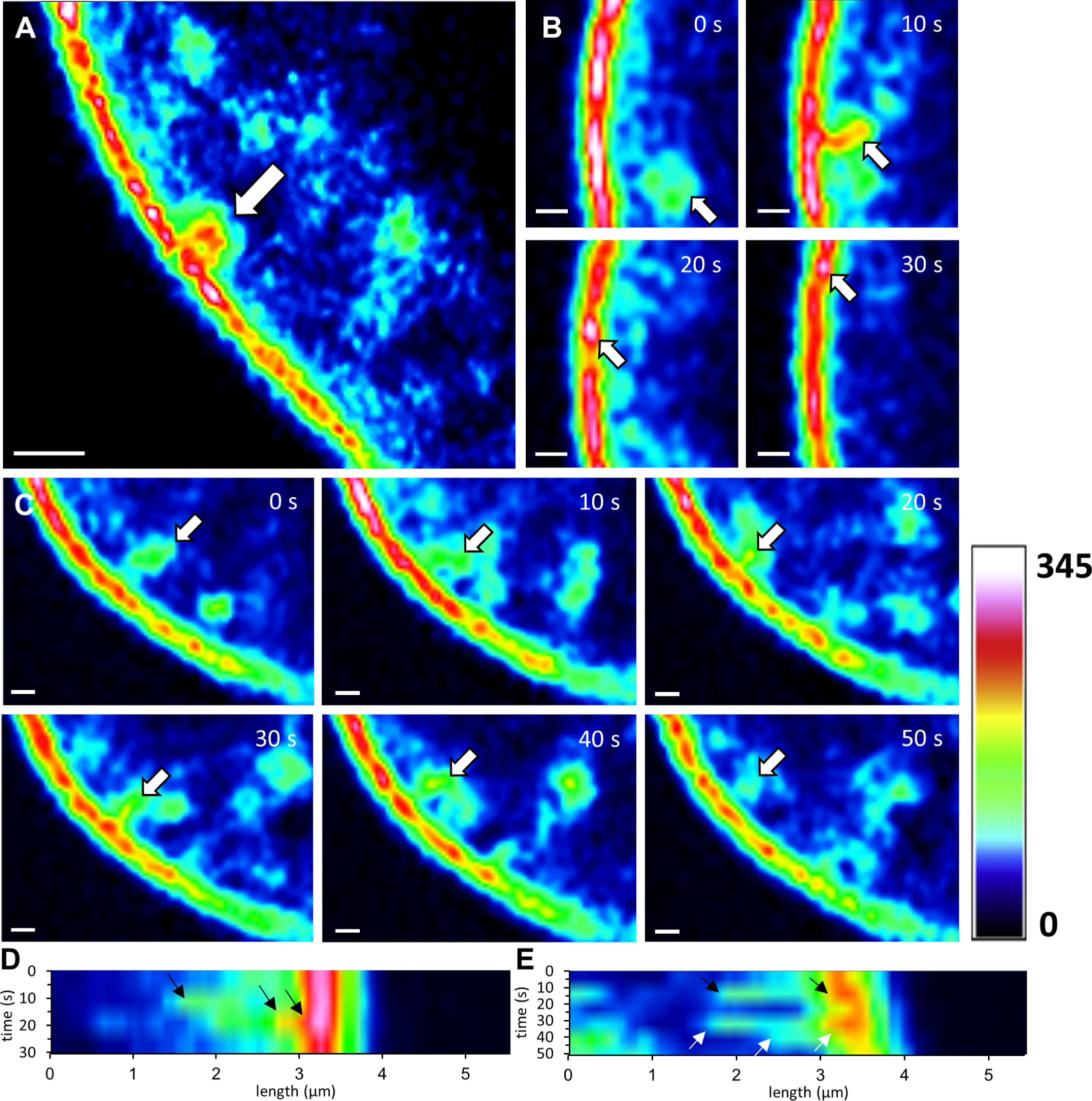
Time-course visualization of close contacts between moving compartments containing GFP-RHD2 and the apical plasma membrane in the root hair apex. **(A)** Round-shaped compartment (arrow) in direct contact with strong GFP-RHD2 fluorescence signal in the apical plasma membrane. **(B, C)** Two events of interactions between the compartments and the apical plasma membrane **(Supplemental Video S7)** showing fast full integration with the plasma membrane (arrows; **B, Supplemental Video S8)** or temporal contact lasting 50 seconds without full integration (fusion) with the plasma membrane (arrows; **C, Supplemental Video S9)**. The time-lapse imaging encompassed 30s **(B)** and 50s **(C)**. The fluorescence distribution is visualized through a semi-quantitative signal intensity using a pseudo-color-coded scale, where dark blue represents minimal intensity (0 arbitrary units) and white represents maximum intensity (345 arbitrary units). **(D, E)** Kymographs showing interactions between the compartments and the apical plasma membrane either by full integration to the membrane (black arrows; **D)** or by approaching and a temporal contact with the plasma membrane (black arrows; **E)**, but without fusion and followed by back movement to the cytoplasm (white arrows; **E)**. Scale bar = 1 µm **(A, B)**, 500 nm **(C)**.

### Movements of early endosomes/TGN and late endosomes in rhd2-1 mutant

Colocalization and common characteristics of compartments containing GFP-RHD2 and early endosomes/TGN but not late endosomes (Figures 4 and 5) raised the question of whether endosomal movements are affected in the *rhd2-1* mutant. To address this, we prepared transgenic *rhd2-1* lines carrying the early endosomal/TGN marker GFP-RabA1d and the late endosomal marker RabF2a-YFP. Since the *rhd2-1* mutant only produced short root hairs that were unable to grow, we performed quantitative tracking of endosomal compartments showing targeted movement to the bulge (red group), moving in the cytoplasm outside of the bulge (blue group), and moving towards the bulge and back (cyan group) during a 60 s imaging period. Quantitative determination of the ratio between individual groups revealed a higher proportion of early endosomes/TGN moving to the bulges in the *rhd2-1* mutant (Figure 8A). However, similar to the compartments moving to the bulge and back (the cyan group), they moved over shorter distances compared to the control line (Figure 8B-D). Characterization of movement parameters of GFP-RabA1d compartments in the control line by the frequency distribution diagram revealed different behaviors of individual groups (Figure 8E and F). In particular, the curves for maximum speed were shifted differentially (Figure 8E). This indicates different properties of early endosomal/TGN compartments that participate in bulge formation. In contrast, frequency distribution analysis of the speed among all three different groups of GFP-RabA1d compartments in the *rhd2-1* mutant showed no differences (Figure 8G and H). These data indicated that the movement of early endosomal/TGN compartments was partially affected in *rhd2-1*.

**Figure 8.**
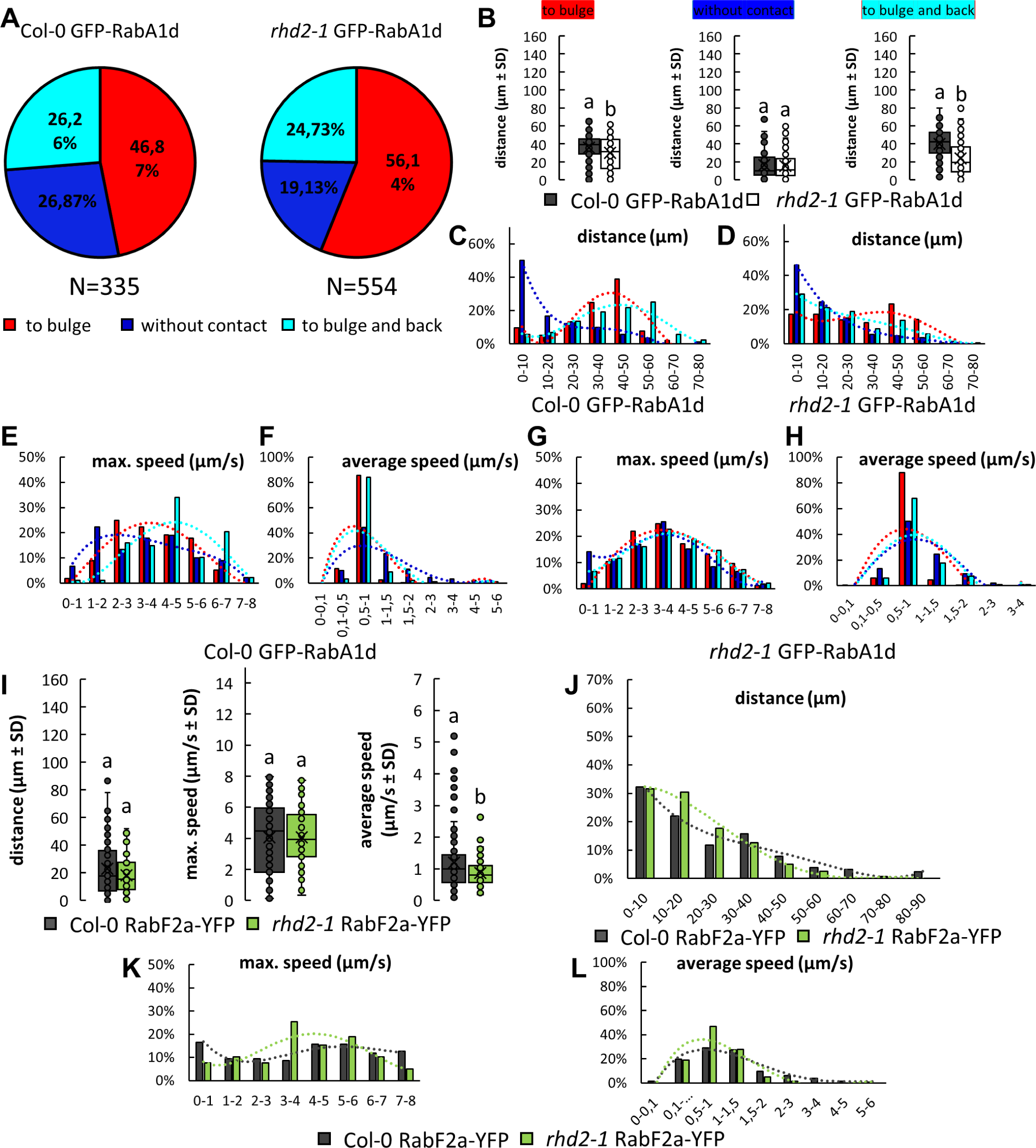
Qualitative and quantitative evaluation of endosomal compartments movements in developing bulges during root hair formation in Arabidopsis control and *rhd2-1* mutant lines. **(A)** Quantitative determination of the ratio between the compartments showing targeted movement to the bulge (red), moving in the cytoplasm outside of the bulge (blue), and moving to the bulge and back (cyan) in the control line (Col-0) and *rhd2-1* mutant bearing early endosomal marker GFP-RabA1d. **(B)** Quantitative evaluation of tracking distance for compartments containing GFP-RabA1d and moving either to the bulge (red), in the cytoplasm outside of the bulge (blue), or to the bulge and back (cyan) in Col-0 control line and *rhd2-1* mutant. **(C, D)** Frequency distributions of distances for tracked compartments moving either to the bulge (red), in the cytoplasm outside of the bulge (blue), or to the bulge and back (cyan) in Col-0 control line **(C)** and *rhd2-1* mutant **(D)** bearing GFP-RabA1d marker. **(E-H)** Frequency distributions of the maximum **(E, G)** and average speed **(F, H)** of the tracked compartments moving either to the bulge (red), in the cytoplasm outside of the bulge (blue), or to the bulge and back (cyan) in Col-0 control line **(E, F)** and *rhd2-1* mutant **(G, H)** carrying GFP-RabA1d. More than 335 compartments from 2 to 4 bulges from individual plants were analyzed. **(I)** Quantitative evaluation of compartment tracking distances, their maximum, and average speed of movement in bulges of Col-0 and *rhd2-1* mutant bearing late endosomal marker RabF2a-YFP. **(J-L)** Frequency distributions of the distance **(J)**, maximum speed **(K)**, and average speed **(L)** of the analyzed compartments moving in bulges of Col-0 control line and *rhd2-1* mutant bearing RabF2a-YFP. More than 127 compartments from 2 to 4 bulges from individual plants were analyzed. Box plots show the first and third quartiles, split by the median (line) and mean (cross). Whiskers extend to include the max/min values. Different lowercase letters indicate statistical significance between treatments (P < 0.05).

Late endosomal compartments did not accumulate in the apex of the growing root hairs (Figure 5B and D; Supplemental Figures S4B and S7B). For this reason, we analyzed all late endosomes carrying RabF2a-YFP within one homogenous group. Characterization of movement distances of RabF2a-YFP compartments in bulging trichoblasts showed only negligible differences between control and *rhd2-1* (Figure 8I and J). In addition, the maximum and average movement speed measurements provided similar values (Figure 8K and L). Thus, the movement of late endosomes seems to be unaffected in *rhd2-1*.

### Distribution of GFP-RHD2 in root hairs after filipin III treatment

The delivery of compartments with GFP-RHD2 to the apical PM in bulges and growing root hairs requires physical interactions with the PM (Figure 7). Simultaneously, we identified and quantitatively characterized a sub-population of GFP-RHD2 compartments moving to and contacting the PM and subsequently moving back to the cytoplasm (Figures 6 and 7). Docking, eventual fusion, or recycling of these compartments back to the cytoplasm may require the participation of integral lipid membrane components. Structural sterols are integral components of plant membranes and are abundant in the apical PM of developing bulges and growing root hairs (Ovečka et al., 2010). In this study, we used filipin III, a vital probe for *in vivo* localization and sequestration of structural sterols (Grebe et al., 2003; Bonneau et al., 2010) in a physiologically high concentration that causes the formation of filipin-sterol complexes at the PM (Milhaud et al., 1998; Bonneau et al., 2010; Ovečka et al., 2010) and a rapid arrest of root hair tip growth (Jones et al., 2006; Ovečka et al., 2010). First, roots of the GFP-RHD2 line were treated with 10 µg.mL^-1^ filipin III using gentle perfusion directly at the microscope stage, and plants were examined by time-lapse live-cell imaging. Growing root hairs were photographed to visualize the cytoarchitecture using differential interference contrast (DIC) optics, while epifluorescence illumination allowed the monitoring of GFP-RHD2 localization and filipin III-induced PM staining during perfusion. The stabilized position of the sample at the microscope stage during the perfusion and sequential image acquisition (Figure 9) and monitoring of GFP-RHD2 accumulation at the tip by measuring fluorescence intensity profiles (Supplemental Figure S12) also provided the means for evaluation of the root hair tip growth. Root hairs before perfusion (time from 0 to 7 min) showed typical cytoarchitecture with tip-focused GFP-RHD2 localization (Figure 9A and B). Analysis of the GFP fluorescence temporal distribution along a median profile in the form of kymograph revealed continuous and sustained tip growth (Figure 9C and D). To avoid any misinterpretations caused by changes induced by mechanical and osmotic disturbances during the application, perfusion with the control medium (mock treatment) was applied to control root hairs (Figure 9A). The control medium application induced a transient movement of the vacuole closer to the tip, slightly reducing the cytoplasm amount in the apical and subapical root hair zone (Figure 9A; DIC, time from 8 to 10 min). It was connected with a transient reduction of GFP-RHD2 fluorescence in the apex (Figure 9A; GFP, time from 8 to 10 min; arrow in Figure 9C). However, both parameters were reversibly restored to normal and maintained until the end of the experiments (Figure 9A; DIC, GFP, time up to 15 min). Thus, after a mock treatment, root hair cytoarchitecture (Figure 9A; DIC, from 11 to 15 min; Supplemental Video S10), tip-focused GFP-RHD2 localization (Figure 9A; GFP, from 11 to 15 min; Supplemental Figure S12A; Supplemental Video S11), and root hair tip growth rate (Figure 9C; from 11 to 15 min) were maintained and not considerably disturbed. As expected, epifluorescence ultra-violet illumination for filipin III excitation produced no signal upon mock treatment (Figure 9A; UV, time from 0 to 15 min).

**Figure 9.**
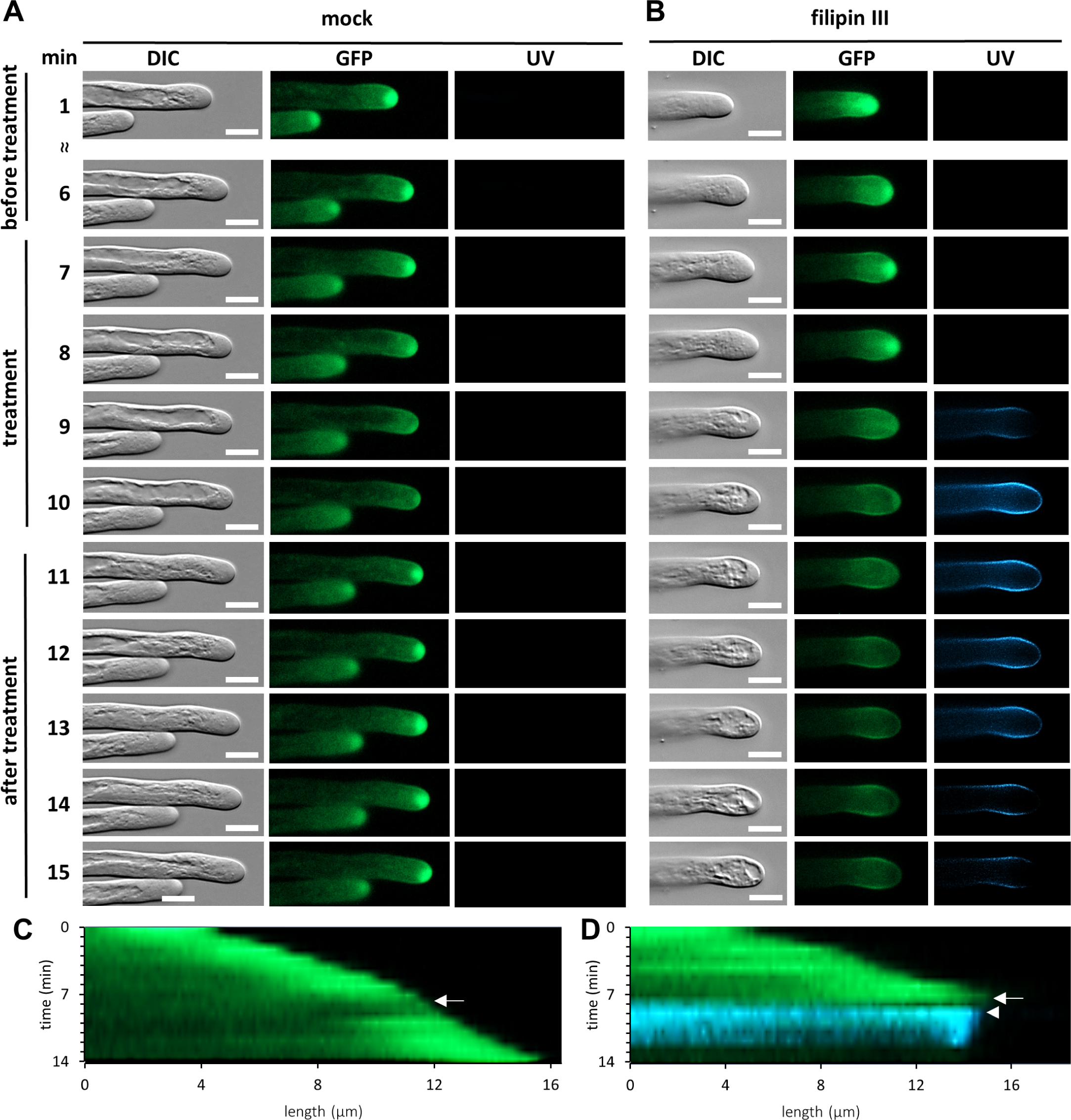
Reaction of growing root hairs and changes of GFP-RHD2 localization after filipin III application. **(A-B)** Growing root hairs imaged either by differential interference contrast (DIC) for cytoarchitecture visualization, blue excitation fluorescence light for GFP-RHD2 localization (GFP), or ultra-violet excitation fluorescence light for visualization of plasma membrane staining by filipin III (UV). Root hairs show normal cytoarchitecture, tip-focused GFP-RHD2 localization, no signal in UV light **(A, B)**, and sustained tip growth **(C, D)** before treatment (time from 0 to 7 min). During perfusion mock treatment **(A)**, there was a transient reduction of the cytoplasm at the subapical root hair zone **(A**; DIC, 8-10 min; **Supplemental Video S10)** and transient reduction of GFP-RHD2 in the apex **(A**; GFP, 9-10 min; **Supplemental Video S11**; arrow in **C)**, while it was reversibly restored and maintained **(B)**, root hairs stopped growth (arrow; **D)**, their tips started to be invaded by a round vacuoles **(B**; DIC, 9-10 min; **Supplemental Video S12)**, and GFP-RHD2 disappeared from the root hair apex **(B**; GFP, 9-10 min; **Supplemental Video S13)**. Simultaneously, filipin III-positive blue signal appeared at the plasma membrane **(B**; UV, 9-10 min; arrowhead in **D)**. After a mock treatment **(A)**, root hair cytoarchitecture **(A**; DIC, 11-15 min), tip-focused GFP-RHD2 localization **(A**; GFP, 11-15 min; **Supplemental Figure S12)**, and root growth rate **(C**; 11-15 min) was regularly maintained and was not disturbed. Root hair after filipin III treatment **(B)** showed disrupted cytoarchitecture such as vacuolated root hair apex **(B**; DIC, 11-15 min), lost GFP-RHD2 from the tip **(B**; GFP, 11-15 min; **Supplemental Figure S12)**, filipin III-positive signal appeared at the plasma membrane **(B**; UV, 11-15 min), and completely blocked tip growth **(D**; 11-15 min). Reduction of filipin III-positive signal at the plasma membrane **(B**; UV, 11-15 min) was caused by a bleaching induced by a wide-field epifluorescence illumination. Scale bar = 10 µm **(A, B)**.

Perfusion with filipin III caused the arrest of root hair tip growth (Figure 9B; DIC, time from 7 to 8 min) and was documented in the related kymograph (Figure 9D, arrow). Consequently, root hair tips were invaded by round articulated vacuoles that irreversibly persisted until the end of the experiments (Figure 9B; DIC, time from 9 to 15 min). As the tip growth was arresting, GFP-RHD2 disappeared irreversibly from the root hair apex (Figure 9B; DIC, time from 9 to 15 min). Alongside the loss of GFP-RHD2, a filipin III-positive blue signal appeared at the plasma membrane (Figure 9B; UV, time from 9 to 15 min; arrowhead in Figure 9D). Therefore, root hairs after filipin III treatment showed disrupted cytoarchitecture due to massive vacuolation of the root hair apex (Figure 9B; DIC, from 9 to 15 min; Supplemental Video S12), a loss of GFP-RHD2 from the tip (Figure 9B; GFP, from 9 to 15 min; Supplemental Figure S12B; Supplemental Video S13), and accumulated filipin III-positive signal at the plasma membrane (Figure 9B; UV, from 9 to 15 min), resulting in blocked tip growth (Figure 9D; from 8 to 15 min).

Next, the root hairs of the GFP-RHD2 line were treated with 10 µg.mL^-1^ filipin III for 10 min, and subcellular changes in GFP-RHD2 distribution were directly examined using a spinning disk microscope. The appearance of filipin III fluorescence in the PM of root hairs after application coincided with the tip growth arrest. Treatment with filipin III led to a layer-like accumulation of GFP-RHD2 closely associated with PM at the apex of root hairs (Figure 10A and C). Because filipin III interacts specifically with sterols in the PM (Milhaud et al., 1998; Bonneau et al., 2010), filipin III-positive blue staining served as a PM marker under these circumstances. Therefore, the layer-like accumulation of GFP-RHD2 beneath the PM suggests that GFP-RHD2 was prevented from being incorporated into the PM after filipin III treatment. Consistent with the localization pattern of GFP-RHD2 in the apical and subapical PM domains in control growing root hairs (Figures 2J-M; 3A and B; 4A and E; 5A and B; 6E and G; Supplemental Figure S1C and E), the extent of layer-like accumulation of GFP-RHD2 after filipin III treatment reached only PM at the apical and subapical domains of root hairs (Figure 10A; Supplemental Figures S13B and S16A). This localization pattern was unique because equivalent experiments on GFP-RabA1d root hairs treated with filipin III revealed accumulation of early endosomal/TGN compartments in the apical part of the root hairs but not in close contact with filipin III-positive apical PM (Figure 10B and E). Semi-quantitative analysis of the fluorescence intensity distribution using a profile function across the root hair apex (Figure 10C) revealed a “basal” level of GFP-RHD2 in filipin III-positive PM but mostly accumulated in a thin layer beneath the PM (Figure 10D). Simultaneously, the clear zone was depleted of GFP-RHD2 fluorescence after filipin III treatment (Figure 10C and D). The same pattern of layer-like accumulation of GFP-RHD2 beneath the PM has been observed in emerged bulges and elongated root hairs (Supplemental Figure S13). Interestingly, GFP-RabA1d accumulation in the clear zone after filipin III treatment showed a different pattern (Figure 10E) because it was detached and did not show direct contact with the filipin III-positive PM (Figure 10E and F). Different GFP-RHD2 and GFP-RabA1d localization patterns after filipin III treatment were repeatedly observed in the apical part of treated root hairs (Supplemental Figure S14). In contrast, treatment of root hairs with filipin III did not cause any changes in the subcellular localization pattern of late endosomes visualized by RabF2a-YFP in root hairs (Supplemental Figure S15). Although root hair tip growth was arrested by filipin III treatment, the distribution of late endosomal compartments carrying RabF2a-YFP resembled the situation under control conditions (Supplemental Figure S4B). Moreover, similar results were obtained in the root hairs of the double GFP-RHD2 × mCherry-RabF2b transgenic line treated with filipin III (Supplemental Figure S16A). GFP-RHD2 accumulated strictly in close association with the apical PM, but late endosomal compartments carrying mCherry-RabF2b were distributed along the whole root hairs (Supplemental Figure S16A). Semi-quantitative fluorescence intensity evaluation at the root hair apex confirmed the close association of GFP-RHD2 with the apical PM and the absence of late endosomal compartments carrying RabF2a-YFP in the clear zone (Supplemental Figure S16B and C). These experiments with filipin III suggest that structural sterols, which are abundant in the apical PM of growing root hairs (Ovečka et al., 2010), might be involved in the dynamic relocation of AtRBOHC/RHD2. Upon filipin III treatment, we did not observe incorporation of GFP-RHD2 into the apical PM and have concluded that structural sterols in PM might be critically involved in AtRBOHC/RHD2 physical incorporation into the apical PM. Based on the fact that PM labeling by filipin III leads to a concentration-dependent formation of filipin-sterol complexes at the ultrastructural level in plant cells (Grebe et al., 2003; Bonneau et al., 2010), including growing root hairs (Ovečka et al., 2010), this treatment leads irreversibly to arrest of the root hair tip growth (Ovečka et al., 2010). These results strongly indicate the dependence of growing root hairs on RHD2/AtRBOHC integration to the apical PM, which requires the participation of structural sterols.

**Figure 10.**
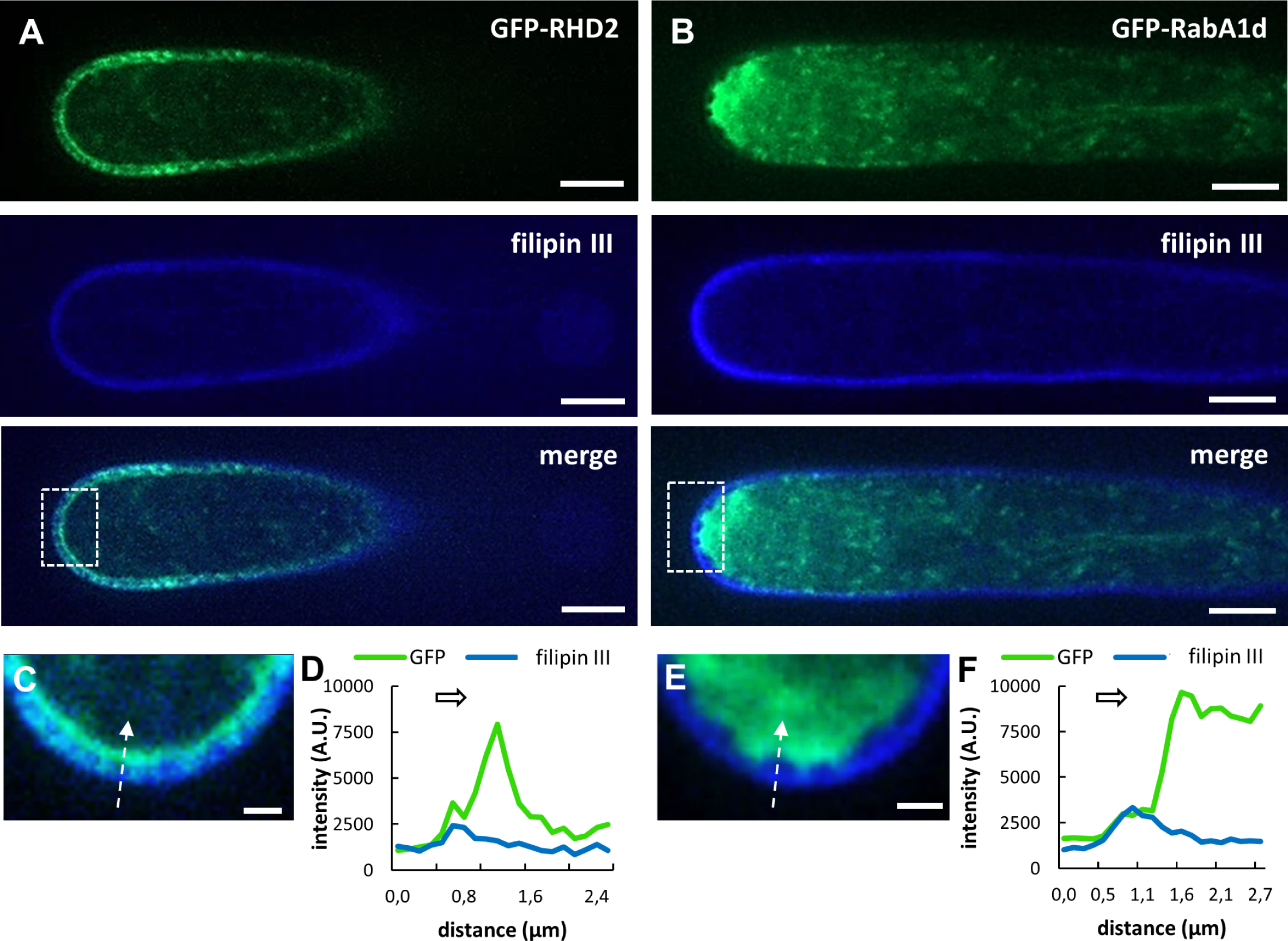
Localization analysis of GFP-RHD2 and GFP-RabA1d in root hairs after treatment with filipin III. **(A)** Localization of GFP-RHD2 in the root hair, filipin III-positive signal (blue) at the plasma membrane, and merged image showing the distribution pattern of GFP-RHD2 after filipin III treatment for 10 min. **(B)** Localization of early endosomal marker GFP-RabA1d in the root hair, filipin III-positive signal at the plasma membrane, and merged image showing the distribution pattern of GFP-RabA1d after filipin III treatment for 10 min. **(C)** Detailed image of the apical part of the root hair within the white box in **(A)**. Arrow indicates the position and the direction of the fluorescence intensity profile measurement. **(D)** Fluorescence intensity profile of the GFP-RHD2 and filipin III-positive signal along the arrow in **(C)**. The arrow above the curves indicates the direction of the measurement. **(E)** Detailed image of the apical part of the root hair within the white box in **(B)**. Arrow indicates the position and the direction of the fluorescence intensity profile measurement. **(F)** Fluorescence intensity profile of the RabA1d-GFP and filipin III-positive signal along the arrow in **(E)**. The arrow above the curves indicates the direction of the measurement. Scale bar = 5 µm **(A, B)**, 1 µm **(C, E)**.

## Discussion

The tip growth of root hairs is a highly polarized process depending on targeted vesicular transport, localized cell wall deposition, and PM extension at the growing tip. It is supported by tip-localized transmembrane gradients (Ca^2+^, ROS, and pH) and by polarized organization and dynamics of the cytoskeleton (Šamaj et al., 2006). The establishment and maintenance of these gradients in root hairs are orchestrated by the NADPH oxidase AtRBOHC/RHD2 (Foreman et al., 2003) activated by Rop GTPases (Jones et al., 2007) accumulated at the site of future root hair initiation (Molendijk et al., 2001; Jones et al., 2002). Loss of AtRBOHC/RHD2 function in *rhd2* mutants leads to the interruption of root hair tip growth. Nevertheless, fully rescued root hair development has been achieved by transforming *rhd2* by GFP-tagged AtRBOHC/RHD2 under the control of the native promoter (Takeda et al., 2008). This also allowed cell-type-specific and developmentally-regulated localization of GFP-RHD2 in trichoblasts, pointing to selective accumulation in bulges and tips of developing root hairs (Takeda et al., 2008). Using different advanced microscopy methods, we qualitatively and quantitatively characterized the subcellular localization of GFP-RHD2 and visualized dynamic vesicular compartments linked with the formation and maintenance of the apical PM domain during root hair development. In root development, these compartments appeared first in the epidermis of the root hair formation zone (the proximal part of the root elongation zone and in the differentiation zone). The number of compartments was significantly higher in the trichoblasts. This localization pattern, spatiotemporally related to root hair formation, is consistent with quantitative microarray analysis of roots, showing enhanced *RHD2* transcription in differentiation and elongation root zones, epidermal cells, and root hairs (Chapman et al., 2019). Next, we used a quantitative particle tracking approach to characterize the temporal and spatial AtRBOHC/RHD2 subcellular redistribution pattern during root hair initiation and growth. This analysis revealed a mode of targeted GFP-RHD2 delivery to the apical PM domain of developing bulges and growing root hairs. Notably, the PM area containing GFP-RHD2 was restricted, and the range of this area correlated with particular stages of root hair development.

Localized delivery of GFP-RHD2 and its incorporation into the PM by motile compartments raises questions about their molecular signatures. As expected, the membrane-specific dye FM4-64 colocalized with GFP-RHD2 at the PM of root hairs in the apical and subapical zones where GFP-RHD2 was restricted. In addition, labeling growing root hairs with FM4-64 revealed a high degree of colocalization with GFP-RHD2 in compartments moving in the cytoplasm of root hairs. This unequivocally indicated their vesicular nature. The short incubation time of growing root hairs with FM4-64 serves to visualize early endocytic compartments (Ovečka et al., 2005). Colocalization studies with early endosomes/TGN (GFP-RabA1d; Ovečka et al., 2010; Berson et al., 2014) or late endosomes (RabF2a-YFP; Voigt et al., 2005; Berson et al., 2014) confirmed that a short FM4-64 application is suitable for the localization of early but not late endosomes. To corroborate these data, we analyzed colocalization in double transgenic lines carrying GFP-RHD2 together with mCherry-VTI12 (an early endosomes/TGN marker; Geldner et al., 2009) and mCherry-RabF2b (a late endosomal marker; Geldner et al., 2009). Using this approach, the colocalization of GFP-RHD2 with the early endosome/TGN marker mCherry-VTI12 was confirmed in the cytoplasmic compartments of growing root hairs. The maximum speed of motile compartments containing GFP-RHD2 in root hairs was typically 4 - 8 μm/s. This is consistent with the dynamic properties of early endosomal compartments carrying molecular markers GFP-RabA1d and YFP-VTI12, with a maximal speed of 6 - 9 μm/s (von Wangenheim et al., 2016).

Importantly, the spatial control of GFP-RHD2 incorporation into the apical PM was closely related to the non-uniform pattern of vesicular movements. The ratio of the compartments moving and fusing to the apical PM, those moving towards, contacting the PM, and subsequently moving back to the cytoplasm, or those moving in the cytoplasm without any contact with the apical PM, changed with different stages of root hair development. The particle tracking analysis provided quantitative parameters of these specific movements and their specific features, especially in compartments coming into contact with the PM. The enlargement of the bulging domain was tightly connected with the higher portion of compartments moving to the apical PM and fusing with it. This correlated with the gradual enlargement of the apical PM area, incorporating and accumulating GFP-RHD2. In contrast, there was an increase in the fraction of compartments contacting the PM and subsequently moving back to the cytoplasm during the growth of root hairs. This likely represents the active recycling of GFP-RHD2 between the PM and motile vesicular compartments in the cytoplasm. These data also indicate that compartments containing GFP-RHD2 are spatiotemporally specialized in particular stages of root hair formation. Consequently, the ratio between different compartment types responsible for the rate of GFP-RHD2 incorporation into the PM and/or recycling was regulated according to the developmental stage of the root hair.

We also tested a possible link between AtRBOHC/RHD2 and the movement of endosomal compartments through detailed quantitative characterization of early endosomes/TGN and late endosomes in the *rhd2-1* mutant. We compared early endosomal/TGN compartments visualized by GFP-RabA1d in the control line, specifically those moving to the bulge or those moving to the bulge and back to the trichoblast and found differences in their speed and trajectory values. The same comparison in the *rhd2-1* mutant transgenic line, however, revealed no differences. It seems that the loss of function of AtRBOHC/RHD2 in the *rhd2-1* mutant influenced, at least partially, the dynamic behavior of early endosomes/TGN in bulges. Comparison with late endosomes visualized by RabF2a-YFP under the same circumstances revealed no differences between the control and *rhd2-1* mutant. These results indicate that *AtRBOHC/RHD2* loss-of-function mutation affects only early endosomal/TGN compartments, verifying the crucial role of early endosomal/TGN compartments in root hair formation and growth. Overall, live-cell localization of GFP-RHD2 during root hair initiation and growth by advanced microscopy indicated the involvement of AtRBOHC/RHD2 in root hair development. Moreover, this approach revealed the involvement of AtRBOHC/RHD2 in the specialization and maintenance of apical PM domains in bulges and tip-growing root hairs as well as targeted delivery to the cortical cytoplasm and the clear zone in dynamic vesicular compartments.

Finally, we tested the role of structural sterols in docking and fusion of compartments containing GFP-RHD2 with the apical PM of growing root hairs. The subcellular localization of structural sterols can be performed using vital fluorescent probes. Filipin III is a polyene antibiotic fluorochrome suitable for the localization of structural sterols in animal (Nichols et al., 2001), yeast (Wachtler et al., 2003), and plant cells (Grebe et al., 2003; Boutté et al., 2009; Liu et al., 2009; Bonneau et al., 2010). Specific accumulation of filipin III signal was observed in the PM of the pre-bulging site, developing bulges, and at the tips of growing root hairs (Ovečka et al., 2010). Filipin III, at higher concentrations or after prolonged exposure, induced the formation of filipin-sterol complexes (Milhaud et al., 1998; Bonneau et al., 2010; Ovečka et al., 2010). This leads to PM damage and changes in physiological functions, and such treatments block root hair tip growth and affect vesicular trafficking (Ovečka et al., 2010). Treatment of root hairs with filipin III allowed as to visualize sterols in the PM of growing root hairs (Figure 9), which was inevitably connected with the arrest of tip growth. Detailed microscopic analysis revealed that filipin III prevented the correct incorporation of GFP-RHD2 into the apical PM zone (Figure 10). Ultrastructural examination documented that concentrations of filipin III at and above 10 µg.mL^-1^ cause the formation of filipin-sterol complexes at the PM (Grebe et al., 2003; Bonneau et al., 2010; Ovečka et al., 2010). The formation of filipin-sterol complexes at the PM was immediately connected with PM rigidification and triggered ion leakage, extracellular pH alkalization, and changes in signaling dependent on protein kinase-mediated phosphorylation (Bonneau et al., 2010). Thus, the formation of filipin III-induced complexes with sterols most likely inhibits sterol functions at the PM, including the proper incorporation of GFP-RHD2 to the apical PM zone of growing root hairs. It was reported previously that treatment with the sterol-disrupting agent methyl-ß-cyclodextrin caused redistribution, clustering, and changes in the diffusion properties of GFP-RBOHD, a fluorescently tagged NADPH oxidase involved in plant defense within the PM of the hypocotyl epidermal cells (Hao et al., 2014). Defective localization of RHD2 NADPH oxidase in root hairs was observed in a sterol-deficient *drought hypersensitive/squalene epoxidase 1-5* (*dry2/sqe1-5*) mutant of *A. thaliana* (Posé et al., 2009; Sena et al., 2017). The link between structural sterols at the PM and root hair formation might be mediated by a regulatory role of Rho of plants (ROPs) such as ROP2, ROP4, and ROP6. ROPs are recruited at the site of future root hair initiation by RopGEF3 (Denninger et al., 2019) and accumulate in bulges and the apex of growing root hairs (Molendijk et al., 2001; Jones et al., 2002). It has been shown that ROP2 and ROP4 activate AtRBOHC/RHD2 for localized ROS production during root hair formation (Carol et al., 2005). ROPs activation leads to transient S-acylation, which is required for their functionality and stabilizes their interaction with the PM and accumulation in lipid rafts (Sorek et al., 2011, 2017). ROP2 and ROP6 colocalize with sterol-enriched PM domains at root hair formation sites and the tips of growing root hairs (Stanislas et al., 2015).

Surprisingly, unlike GFP-RabA1d, GFP-RHD2 in filipin III-treated root hairs accumulated within a thin layer closely associated with the inner cytoplasmic side of the apical PM, and the GFP-RHD2 signal was depleted in the clear zone. These data suggest that structural sterols, which are responsible for membrane fluidity and the formation of lipid nanodomains, are indispensable for the correct insertion of AtRBOHC/RHD2 to the apical PM in root hairs. Such layer-like accumulation of GFP-RHD2 beneath the apical PM suggests that the physical incorporation is prevented, and the depletion of GFP-RHD2 compartments from the clear zone indicates that recycling from the PM was inhibited. This observation concludes that structural sterols seem to play important roles in localized anchoring and maintenance of GFP-RHD2 in the apical PM zone of developing root hairs.

## Material and methods

### Plant material preparation

Experiments were performed with plants of *Arabidopsis thaliana*, wild-type of the Col-0 ecotype, *rhd2-1* (*root hair defective 2*; Foreman et al., 2003) mutant, and stably transformed *A. thaliana* plants carrying constructs with fluorescent markers of endosomal compartments. Stable transgenic lines in the Col-0 and *rhd2-1* backgrounds, bearing fluorescent markers of endosomal compartments, were prepared by crossing. Plants of the *rhd2-1* mutant, used as female donors, were crossed with transgenic plants bearing fluorescent markers for early endosomes/TGN, GFP-RabA1d (Ovečka et al., 2010) or late endosomes, RabF2a-YFP (Voigt et al., 2005). Transgenic plants bearing fluorescent markers of endosomal compartments were used as male donors. Transgenic plants with N-terminal fusion of RHD2 with GREEN FLUORESCENT PROTEIN (GFP) under the control of the *RHD2* promoter (GFP-RHD2 line, cloned and plants prepared by Takeda et al., 2008), complementing the *rhd2* homozygous mutant phenotype (GABI-Kat line 841E09 with T-DNA inserted in the fifth exon; Takeda et al., 2008), were crossed with transgenic plants carrying fluorescent markers of endosomal compartments (Wave line W13R producing mCherry-VTI12 for early endosomes/TGN; Geldner et al., 2009, and Wave line W2R producing mCherry-RabF2b for late endosomes; Geldner et al., 2009). Transgenic plants with GFP-RHD2 were used as female donors in crossings. The progeny of the F1 generation was selected according to fluorescence signals using an Axio Zoom.V16 fluorescence stereomicroscope (Carl Zeiss, Germany), and plants of the F2 or F3 generations were used for experiments. We analyzed the transmission of the *GFP-RHD2/rhd2* mutant and red marker alleles in the F2 and F3 generations by phenotypical analysis. As expected, the progeny of heterozygous plants consisted of approximately 50% of the individuals carrying GFP and red fluorescence simultaneously. GFP fluorescence alone was detected in approximately 25% of plants, analogous to mCherry (Tab. 1). Lines showing both types of fluorescence with the wild-type root hair phenotype were used further for the colocalization studies.

**Table 1.** F2 progeny after crossing of the complemented mutant (*GFP-RHD2/rhd2*) with transgenic lines bearing mCherry-tagged endosomal markers.

| Crossed lines | Phenotype (fluorescence) |  |  |  | No of plants |
| --- | --- | --- | --- | --- | --- |
|  | GFP+mCherry | GFP | mCherry | None |  |
| *GFP-RHD2/rhd2* × *mCherry-RabF2b* | 49% | 25% | 21% | 5% | 118 |
| *GFP-RHD2/rhd2* × *mCherry-VTI12* | 55% | 21% | 17% | 6% | 94 |

### In vitro cultivation

Seeds of *A. thaliana* control, mutant, and transgenic lines were surface-sterilized and planted on ½ MS medium (Murashige and Skoog, 1962) without vitamins, solidified with 0.6% Gellan gum (Alfa Aesar, ThermoFisher). Petri dishes with seeds were stratified at 4 °C for 3 days for synchronized germination. After stratification, seeds were cultivated vertically in environmental chamber at 21 °C, 70% humidity, and 16/8 h light/dark cycle. The illumination intensity was 130 μmol m^-2^s^-1^.

### Light-sheet fluorescence microscopy

Samples for light-sheet fluorescence microscopy (LSFM) were prepared as described by Ovečka et al. (2015). Seeds of GFP-RHD2 line were surface sterilized and transferred to round 90 mm × 25 mm Petri dishes filled with 80 mL of modified MS medium (Ovečka et al. 2014) at pH 5.7, solidified with 0.5% (w/v) Gellan gum (Alfa Aesar, ThermoFisher). After germination, the roots began to grow into the medium gravitropically. Three-days-old plants were inserted into the fluorinated ethylene propylene (FEP) tubes with the medium surrounding the root (root was embedded in the medium, and the upper green part was exposed to air). The samples were transferred to a pre-tempered (22 °C) observation chamber of the LSFM filled with a modified liquid MS medium. The agar block with the plant was partially pushed out from the FEP tube into the liquid medium for imaging of the root tip not surrounded by FEP tube walls. After 30 min of stabilization, plants were observed using a light-sheet Z.1 fluorescence microscope (Carl Zeiss, Germany) equipped with two LSFM 10×/0.2 NA illumination objectives (Carl Zeiss, Germany), Plan-Apochromat 20×/1.0 NA water immersion detection objective (Carl Zeiss, Germany), and the PCO Edge sCMOS camera (PCO AG, Germany) with an exposure time of 40 ms. Imaging was carried out using dual-side illumination and pivot scan mode with a light-sheet thickness of 4.52 μm. GFP was excited at 488 nm and detected between 505 and 545 nm. Images were acquired in three subsequent views aligned to each other in the root growth direction (along the y coordinate) at time points of every 5 min in Z-stack mode for 15 h. Image scaling in x, y, z was adjusted to 0.228 μm × 0.228 μm × 0.477 μm.

### Airyscan confocal laser scanning microscopy (Airyscan CLSM) and spinning disk microscopy (SD)

Plants 2 - 3 days old were transferred to microscopic chambers containing liquid modified MS medium according to Ovečka et al. (2005). After manipulation with plants during sample preparation, the subsequent stabilization period for 24 h allowed undisturbed growth of the root and the formation of new root hairs. The development of root hairs was observed using a confocal laser scanning microscope LSM880 equipped with Airyscan (Carl Zeiss, Germany) and a Cell Observer SD Axio Observer Z1 spinning disk microscope (Carl Zeiss, Germany). Airyscan CLSM was used for qualitative and quantitative characterization of GFP-RHD2 positive compartments during root hair development and growth. Image acquisition was performed with a 20×/0.8 NA dry Plan-Apochromat objective (Carl Zeiss, Germany). The samples were imaged with an excitation laser line of 488 nm and BP420-480 + BP495-550 emission filters for GFP detection. The laser power did not exceed 0.8% of the available laser intensity range. The samples were scanned every 706 ms with 700 ms of exposure time using a 32 GaAsP detector. A spinning disk microscope equipped with alpha Plan-Apochromat 100x /1.57 NA DIC Korr oil immersion objective (Carl Zeiss, Germany) was used for colocalization analyses. The samples were imaged using an excitation laser line of 405 nm and emission filter BP450/50 for filipin III signal, excitation laser line of 488 nm and emission filter BP525/50 for GFP signal, excitation laser line of 514 nm and emission filter BP535/30 for YFP signal, excitation laser line of 514 nm and emission filter BP690/50 for FM4-64 signal, and excitation laser line of 561 nm and emission filter BP629/62 for the mCherry signal. Images were acquired sequentially or simultaneously with two Evolve 512 EMCCD cameras (Photometrics) with an exposure time of 700 ms. The samples were scanned every 731 ms using a camera-streaming mode.

### FM4-64 staining and filipin III application

FM4-64 (Thermo Fisher Scientific) was used as the PM and vesicular marker in the root hairs. Filipin III (Sigma-Aldrich, USA) was used as a vital probe for PM structural sterols, allowing their cross-linking in root hairs at higher concentrations. FM4-64 (4 µmol.L^-1^) and filipin III (10 µg.mL^-1^) in modified liquid MS medium individually or in a mixture were applied by perfusion directly to the microscopic chamber. The total volume of the modified liquid MS medium with FM4-64 or filipin III applied was 100 µL, added in 10 separate steps of 10 µL each for 10 min. After perfusion, the plants were directly observed using a spinning disk microscope.

### Life imaging of GFP-RHD2 in root hairs after filipin III application

Plants, 2 - 3 days old, were transferred to microscopy chambers containing liquid modified MS medium according to Ovečka et al. (2005). Subsequent stabilization for 24 h in a glass cuvette in an environmental chamber allowed the continuation of root growth and the formation of new root hairs. Growing root hairs (n = 3 - 5 root hairs) were observed using an epifluorescence microscope Zeiss Axio Imager M2 equipped with DIC optics and epifluorescence metal halide source illuminator HXP 120 V (Zeiss, Oberkochen, Germany), and analyzed using Zeiss ZEN 2012 Blue software (Zeiss, Germany). Root hairs were recorded at 1 min intervals for 5 min before the application, then for another 5 min throughout the application of mock and 10 μg.mL^-1^ filipin III in modified liquid MS medium, respectively, and 5 min after application. The total volume of applied solutions (both control medium and filipin III) was 50 μL, added sequentially in 5 separate steps of 10 μL each. Gentle application by perfusion was performed directly at the microscope stage in the sample chambers at the respective time points of acquisition. Imaging was performed with a Plan-Neofluar 40×/0.75 NA dry objective and documented with a Zeiss AxioCam ICm1 camera. To visualize the GFP signal, a filter set providing a wavelength of 450 - 490 nm for the excitation and 515 - 565 nm for the emission at 400 ms exposure time was used. For the filipin III signal, the filter set provided an excitation wavelength of 335 - 383 nm and an emission wavelength of 420 - 470 nm was used. The exposure time was 200 ms for both filipin III and DIC channel acquisitions.

### Colocalization analysis

The colocalization of fluorescence signals was analyzed using a spinning disk microscope by simultaneous signal acquisition with two independent Evolve 512 EMCCD cameras. After camera calibration for proper alignment, the fluorescence signals of the two markers were imaged and recorded using a camera streaming mode. Semi-quantitative signal intensity analysis was performed on one selected Z-stack of the scanned area. The signal intensity and mode of colocalization were analyzed in double transgenic lines GFP-RHD2 × mCherry-VTI12 and GFP-RHD2 × mCherry-RabF2b. Similarly, the fluorescence intensity was determined in single transgenic lines GFP-RHD2, GFP-RabA1d, and RabF2a-YFP stained with FM4-64. Profiles for quantitative signal intensity distribution and colocalization were generated using Zen Blue 2014 software (Carl Zeiss, Germany) and graphically edited in Microsoft Excel. Quantitative colocalization analysis of vesicular compartments was conducted within selected ROIs from five individual root hairs at five different time points. Three ROIs selected at each analyzed time point led to 75 measurements of vesicular compartments performed in total. Analogously, colocalization analysis was performed on plasma membranes from five individual root hairs at five different time points. A particular ROI covering the membrane area of each respective time point was selected, providing 25 measurements on plasma membranes in total. Selected ROIs of vesicles and membranes of double lines (GFP-RHD2 × mCherry-VTI12 and GFP-RHD2 × mCherry-RabF2b) and FM4-64 staining of GFP-RHD2, GFP-RabA1d, and RabF2a-YFP lines were subjected to quantitative colocalization analysis using Pearsońs correlation coefficients in Zen Blue 2014 software (Carl Zeiss, Germany). The results were graphically edited using Microsoft Excel.

### 3D rendering of GFP-RHD2 distribution

Data obtained by LSFM time-lapse imaging from growing primary roots and root hairs of the GFP-RHD2 line were subjected to 3D rendering. In Zen 2014 software, black edition (Carl Zeiss, Germany), a subset of selected time-points (lasting for 4 h and 25 min of imaging) was created from the whole volume of the root (containing 483 Z-stacks). The subsets were selected to capture the different developmental stages of root hair formation in the root differentiation zone. The output data were imported into Arivis Vision4D 2.12.6 software (Arivis AG, Germany) and automatically converted to the *.sis file. Subsequently, data were visualized as 3-D objects by activation of the 4-D viewer in the Viewer Types Panel. 3-D objects were rendered in the maximum intensity mode. The color scale was switched to rainbow coloring to enhance the contrast visibility of the fluorescence signal intensities and distribution. The lowest and highest saturation points were adjusted to 219 (black) and 427 (red), respectively. For 3-D visualizations of primary root cross-sections, clipping around a region of interest was adjusted using a 4-D clipping panel and ROI tab. Videos were configured by arranging keyframes and rendered as movies for export in the storyboard extension. Animations were prepared by clipping the 3-D model against the y- and z-planes in a 4-D clipping panel and using rotation and zoom tools.

### Single-particle tracking of compartments containing GFP-RHD2

Arivis Vision4D program, version 3.1.4. (Arivis AG, Germany) was used for quantitative tracking of the GFP-RHD2 compartments. Three-days-old plants were imaged using Airyscan CLSM. Different stages of root hair development were selected, and one particular Z-stack of the field of view was imaged for 60 s. Acquired data were processed and quantified using the Tracked cells function of the Arivis software. Before analysis, the following conditions were set up for normalization: (i) the highest and lowest fluorescence intensity thresholds were set to 30 and 410, respectively, (ii) the distance of compartment movement recorded during one time period should not exceed 5 µm, (iii) accepted compartments did not disappear from the image plane for more than two time periods during acquisition, and (iv) the accepted compartment surface area was always in the range of 0.5 - 3 µm^2^. Characterized parameters were minimum, maximum, and average speed, the distance of GFP-RHD2 compartments moving in different stages of root hair development, namely in the early bulge, late bulge, short growing root hairs, and longer growing root hairs. The analyzed compartments were divided into the following subgroups: moving to and contacting with the apical PM, in the cytoplasm without any contact with the PM, and contacting the PM and moving back to the cytoplasm. A comparison was made between compartments moving outside of the bulging domain in trichoblasts and those moving in atrichoblasts. The individual color-coded trajectories of the analyzed compartments were displayed directly in the images. Arivis software was also used to generate the 2.5-D fluorescence intensity profiles.

### Data analysis and graphical presentation

Kymographs were generated from time-lapsed images acquired by LSFM, Airyscan CLSM, and SD using the appropriate plugin of Zen software (Blue version). All graphs were prepared using Microsoft Excel software. Statistical significance (p < 0.05) was determined using the STATISTICA 12 (StatSoft, TIBCO Software Inc., Palo Alto, CA, USA) by ANOVA and subsequent Fisher’s LSD test (p < 0.05).

## Acknowledgments

We thank Liam Dolan for the seeds of the *rhd2-1* mutant and rescued the *GFP-RHD2/rhd2* line. We would like to thank Editage (www.editage.com) for English language editing.

## Supplemental Materials

**Supplemental Figure S1. Semi-quantitative analysis of root hair tip growth rate in individual developmental stages. (A, C, E, G)** Representative images from time-lapse recordings of root hairs using Airyscan confocal laser scanning microscopy. **(B, D, F, H)** Semi-quantitative analysis of root hair growth rates presented by kymographs made along dotted white lines in **(A, C, E, G)**, encompassing the distances of 30 - 40 µm and the time periods of 8 - 16 min. Measured stages are the bulge **(A, B)**, short root hair **(C, D)**, long root hair **(E, F)**, and non-growing root hair **(G, H)**. Scale bar = 5 µm **(A, C, E, G)**.

**Supplemental Figure S2. Semi-quantitative analysis of the tip-focused GFP-RHD2 spatial signal intensity distribution obtained using the 2.5-D rendering function in different stages of root hair development. (A)** early bulge, **(B)** late bulge, **(C)** short, and **(D)** longer growing root hairs.

**Supplemental Figure S3. Example of a quantitative colocalization analysis between FM4-64 and GFP-RHD2. (A)** Scatter plot showing the colocalization and **(B)** ROIs of selected intracellular compartments used for the colocalization analysis. **(C)** Scatter plot showing the colocalization and **(D)** ROI of selected subapical plasma membrane for the colocalization analysis. Horizontal axis represents GFP channel and vertical axis represents FM4-64 channel. Crossed hairline corresponds to threshold setting according to Costes et al. (2004).

**Supplemental Figure S4. Colocalization analysis of early and late endosomal markers with selective membrane styryl dye FM4-64 in growing root hairs. (A)** Localization pattern of the early endosomal marker GFP-RabA1d in the growing root hair co-labelled with membrane styryl dye FM4-64. In merged image, an extent of fluorescence intensity colocalization has been analysed along the cross-section profile marked by interrupted white line (white arrows indicate the plasma membrane) and in moving compartments depicted by the white box. **(B)** Localization pattern of the late endosomal marker RabF2a-YFP in the growing root hair co-labelled with membrane styryl dye FM4-64. In merged image, an extent of fluorescence intensity colocalization has been analysed along the cross-section profile marked by interrupted white line (white arrows indicate the plasma membrane) and in moving compartments within the white box. **(C, D)** Fluorescence intensity profiles of the GFP-RabA1d and FM4-64 signals **(C)** along the cross-section interrupted white line in **(A)** and fluorescence intensity profiles of the RabF2a-YFP and FM4-64 signals **(D)** along the cross-section interrupted white line in **(B)**. Arrows indicate position of the plasma membrane. **(E,G)** Detailed image of compartments containing GFP-RabA1d and FM4-64 **(E)** within the white box in **(A)** and detailed image of compartments containing RabF2a-YFP and FM4-64 **(G)** within the white box in **(B)**. Arrows indicate positions of the compartments and interrupted white lines depict the profile for fluorescence intensity quantification. **(F, H)** Fluorescence intensity profile of the RabA1d-GFP and FM4-64 signals **(F)** along the interrupted white line in **(E)** and fluorescence intensity profile of the RabF2a-YFP and FM4-64 signal **(H)** along the interrupted white line in **(G)**. Arrows indicate position of the compartments. Scale bar = 5 µm **(A, B)**, 1 µm **(E, G)**.

**Supplemental Figure S5. Example of a quantitative colocalization analysis** between **FM4-64 and early endosomal marker GFP-RabA1d. (A)** Scatter plot showing the colocalization and **(B)** ROIs of selected intracellular compartments used for the colocalization analysis. **(C)** Scatter plot showing the colocalization and **(D)** ROI of selected subapical plasma membrane for the colocalization analysis. Horizontal axis represents GFP channel and vertical axis represents FM4-64 channel. Crossed hairline corresponds to threshold setting according to Costes et al. (2004).

**Supplemental Figure S6. Example of a quantitative colocalization analysis between FM4-64 and late endosomal marker RabF2a-YFP. (A)** Scatter plot showing the colocalization and **(B)** ROIs of selected intracellular compartments used for the colocalization analysis. **(C)**Scatter plot showing the colocalization and **(D)** ROI of selected subapical plasma membrane for the colocalization analysis. Horizontal axis represents GFP channel and vertical axis represents FM4-64 channel. Crossed hairline corresponds to threshold setting according to Costes et al. (2004).

**Supplemental Figure S7. Localization of early and late endosomal markers in growing root hairs of control lines. (A)** Localization of the early endosomal/trans-Golgi marker mCherry-VTI12 in control (Col-0) line. **(B)** Localization of the late endosomal marker mCherry-RabF2b in control (Col-0) line. Scale bar = 5 µm **(A, B)**.

**Supplemental Figure S8. Example of a quantitative colocalization analysis between GFP-RHD2 and early endosomal/trans-Golgi network marker mCherry-VTI12. (A)** Scatter plot showing the colocalization and **(B)** ROIs of selected intracellular compartments used for the colocalization analysis. **(C)** Scatter plot showing the colocalization and **(D)** ROI of selected subapical plasma membrane for the colocalization analysis. Horizontal axis represents GFP channel and vertical axis represents mCherry channel. Crossed hairline corresponds to threshold setting according to Costes et al. (2004).

**Supplemental Figure S9. Example of a semi-quantitative colocalization analysis between GFP-RHD2 and late endosomal marker mCherry-RabF2b. (A)** Scatter plot showing the colocalization and **(B)** ROIs of selected intracellular compartments used for the colocalization analysis. **(C)**Scatter plot showing the colocalization and **(D)** ROI of selected subapical plasma membrane for the colocalization analysis. Horizontal axis represents GFP channel and vertical axis represents mCherry channel. Crossed hairline corresponds to threshold setting according to Costes et al. (2004).

**Supplemental Figure S10. Quantitative evaluation of the tracking parameters of compartments containing GFP-RHD2 in different stages of root hair development.** Distributions of distance frequencies, maximum, and average speeds of analysed compartments moving to either the apical plasma membrane (red), in the cytoplasm without any contact with the apical plasma membrane (blue), or contacting the plasma membrane and moving back to the cytoplasm (cyan) in early bulge, late bulge, short, and longer root hairs. More than 59 compartments from 3 to 7 growing root hairs from individual plants were analyzed.

**Supplemental Figure S11. Qualitative determination and quantitative tracking of moving compartments containing GFP-RHD2 in atrichoblasts and trichoblasts outside of the bulging zone. (A, B)** Sequential line tracking of the compartments with GFP-RHD2 in atrichoblast **(A)** and trichoblast outside of the bulging zone **(B)** in the time period of 60s. **(C,D)** Quantitative determination of compartments tracking distances, minimum, maximum, and average speeds of their movements, as compared separately for atrichoblasts **(C)** and trichoblasts outside of the bulging zone **(D)**. **(E-G)** Distribution of the frequencies of the distance **(E)**, maximum speed **(F)**, and average speed **(G)** of the analysed compartments moving in atrichoblasts and in trichoblasts outside of the bulging zone. More than 105 compartments from 2 to 4 cells from individual plants were analyzed. Box plots show the first and third quartiles, split by the median (line) and mean (cross). Whiskers extend to include the max/min values. Different lowercase letters indicate statistical significance between treatments (P < 0.05). Scale bar = 5 µm (A, B).

**Supplemental Figure S12. Changes of GFP-RHD2 localization in growing root hairs after filipin III application. (A, B)** Growing root hairs imaged using blue excitation fluorescence light for GFP-RHD2 localization and treated by a mock **(A)** or filipin III **(B)**. Distribution of the GFP-RHD2 was analysed by a fluorescence intensity profile measurement along white dotted lines. Fluorescence intensity profiles are shown for every min of time-course acquisition before treatment (0-6 min), during treatment with mock **(A**; 7-10 min) and filipin III **(B**; 7-10 min), and after treatments (11-15 min). Displacement of the fluorescence intensity peak between time points **(A)** corresponds to the root hair tip growth rate, which is not evident during and after filipin III treatment **(B)** since the growth was stopped.

**Supplemental Figure S13. Localization analysis of GFP-RHD2 in bulge and root hair after treatment with filipin III. (A, B)** Localization of GFP-RHD2 in the bulge **(A)** and root hair **(B)**, filipin III-positive signal at the plasma membrane, and merged images showing the distribution pattern of GFP-RHD2 compartments after filipin III treatment for 10 min. **(C,**BOHC/RHD2 physical incorporation into the **E)** Detailed image of the apical part of the bulge **(C)** and root hair **(E)** within the white boxes in **(A)** and **(B)**. Arrows indicate the position and the direction of the fluorescence intensity profile measurement. **(D, F)** Fluorescence intensity profiles of the GFP-RHD2 and filipin III-positive signal along arrows in **(C)** and **(E)**. Arrows above the curves indicate the direction of the measurement. Scale bar = 5 µm **(A, B)**, 1 µm **(C, E)**.

**Supplemental Figure S14. Localization analysis of GFP-RHD2 and** RabA1d**-GFP in root hairs after treatment with filipin III. (A)** Detailed image of the apical part of the root hair showing localization of GFP-RHD2 after filipin III treatment for 10 min. **(B)** Detailed image of the apical part of the root hair depicting localization of RabA1d-GFP after filipin III treatment for 10 min. Five different arrows indicate positions and directions of independent fluorescence intensity profiles which were measured. Scale bar = 1 µm **(A, B)**.

**Supplemental Figure S15. Localization analysis of RabF2a-YFP in** root **hairs after treatment with filipin III. (A)** Localization of RabF2a-YFP in the root hair, filipin III-positive signal at the plasma membrane, and merged image showing the distribution pattern of RabF2a-YFP compartments after filipin III treatment for 10 min. **(B)** Detailed image of the apical part of the root hair within the white box in **(A)**. Arrow indicates the position and the direction of the fluorescence intensity profile measurement. **(C)** Fluorescence intensity profile of the RabF2a-YFP and filipin III-positive signal along the arrow in **(B)**. Arrow above the curves indicates the direction of the measurement. Scale bar = 5 µm **(A)**, 1 µm **(B)**.

**Supplemental Figure S16. Localization analysis of GFP-RHD2 and mCherry-RabF2b in root hairs after treatment with filipin III. (A)** Localization patterns of GFP-RHD2 and late endosomal marker mCherry-RabF2b in the root hair, filipin III-positive signal at the plasma membrane, and merged image showing the distribution of GFP-RHD2 and mCherry-RabF2b after filipin III treatment for 10 min. **(B)** Detailed image of the apical part of the root hair within the white box in **(A)**. Arrow indicates the position and the direction of the fluorescence intensity profile measurement. **(C)** Fluorescence intensity profile of the GFP-RHD2, mCherry-RabF2b and filipin III along the arrow in **(B)**. Arrow above the curves indicates the direction of the measurement. Scale bar = 5 µm **(A)**, 1 µm **(B)**.

**Supplemental Video S1.** 3-D maximum intensity projection and a volumetric rendering of a pseudo-color-coded semi-quantitative fluorescence intensity distribution of GFP-RHD2 in the root hair formation zone in Arabidopsis from light-sheet fluorescence microscopy imaging.

**Supplemental Video S2.** Time-lapse imaging of developing root hair in root of transgenic Arabidopsis plant carrying GFP-RHD2 using light-sheet fluorescence microscopy. Exposure time 40 ms, time point every 5 min, imaging time 220 min.

**Supplemental Video S3.** Time-lapse imaging of developing late bulge using Airyscan confocal laser scanning microscopy showing polarized accumulation of GFP-RHD2 at the apical plasma membrane domain and cortical cytoplasm in a pseudo-color-coded semi-quantitative fluorescence intensity distribution. Exposure time 700 ms, time point every 9.8 s, imaging time 16 min.

**Supplemental Video S4.** Time-lapse imaging of short growing root hair using Airyscan confocal laser scanning microscopy showing polarized accumulation of GFP-RHD2 at the apical plasma membrane domain and cortical cytoplasm in a pseudo-color-coded semi-quantitative fluorescence intensity distribution. Exposure time 700 ms, time point every 9.8 s, imaging time 8 min.

**Supplemental Video S5.** Time-lapse imaging of long growing root hair using Airyscan confocal laser scanning microscopy showing polarized accumulation of GFP-RHD2 at the apical plasma membrane domain and cortical cytoplasm in a pseudo-color-coded semi-quantitative fluorescence intensity distribution. Exposure time 700 ms, time point every 9.8 s, imaging time 8 min.

**Supplemental Video S6.** Time-lapse imaging of non-growing root hair using Airyscan confocal laser scanning microscopy showing polarized accumulation of GFP-RHD2 at the apical plasma membrane domain and cortical cytoplasm in a pseudo-color-coded semi-quantitative fluorescence intensity distribution. Exposure time 700 ms, time point every 9.8 s, imaging time 3 min.

**Supplemental Video S7.** Time-lapse imaging of close contacts (arrows) between moving compartments containing GFP-RHD2 and the apical plasma membrane in the apex of late bulge using Airyscan confocal laser scanning microscopy in a pseudo-color-coded semi-quantitative fluorescence intensity distribution. Exposure time 700 ms, time point every 9.8 s, imaging time 60 s.

**Supplemental Video S8.** Time-lapse imaging of contact between moving compartment containing GFP-RHD2 and the apical plasma membrane in the apex of late bulge showing fast full integration. Imaging using Airyscan confocal laser scanning microscopy in a pseudo-color-coded semi-quantitative fluorescence intensity distribution. Exposure time 700 ms, time point every 9.8 s, imaging time 30 s.

**Supplemental Video S9.** Time-lapse imaging of contact between moving compartment containing GFP-RHD2 and the apical plasma membrane in the apex of late bulge showing temporal contact without full **integration**. Imaging using Airyscan confocal laser scanning microscopy in a pseudo-color-coded semi-quantitative fluorescence intensity distribution. Exposure time 700 ms, time point every 9.8 s, imaging time 50 s.

**Supplemental Video S10.** Time-lapse imaging of growing root hair using DIC optics showing growth rate and cytoarchitecture during perfusion application of control medium. Exposure time 200 ms, time point **every** 1 min, imaging time 14 min.

**Supplemental Video S11.** Time-lapse imaging of growing root hair using fluorescence excitation for localization of GFP-RHD2 during perfusion application of control medium. Exposure time 400 ms, time point every 1 **min**, imaging time 14 min.

**Supplemental Video S12.** Time-lapse imaging of growing root hair using DIC optics showing growth rate and cytoarchitecture during perfusion application of filipin III. Exposure time 200 ms, time point every 1 min, imaging time 14 min.

**Supplemental Video S13.** Time-lapse imaging of growing root hair using fluorescence excitation for localization of GFP-RHD2 during perfusion application of filipin III. Exposure time 400 ms, time point every 1 min, imaging **time** 14 min.

## References

1. Berson T, Wangenheim D, Takáč T, Šamajová O, Rosero A, Ovečka M, Komis G, Stelzer EH, Šamaj J (2014) Trans-Golgi network localized small GTPase RabA1d is involved in cell plate formation and oscillatory root hair growth. BMC Plant Biol 14: 252

2. Bonneau L, Gerbeau-Pissot P, Thomas D, Der C, Lherminier J, Bourque S, Roche Y, Simon-Plas F (2010) Plasma membrane sterol complexation, generated by filipin, triggers signalling responses in tobacco cells. Biochim Biophys Acta 1798: 2150–2159

3. Bottanelli F, Foresti O, Hanton S, Denecke J (2011) Vacuolar transport in tobacco leaf epidermis cells involves a single route for soluble cargo and multiple routes for membrane cargo. Plant Cell 23: 3007–3025

4. Boutté Y, Frescatada-Rosa M, Men S, Chow C-M, Ebine K, Gustavsson A, Johansson L, Ueda T, Moore I, Jürgens G, et al (2009) Endocytosis restricts Arabidopsis KNOLLE syntaxin to the cell division plane during late cytokinesis. EMBO J 29: 546–558

5. Campanoni P, Blatt MR (2007) Membrane trafficking and polar growth in root hairs and pollen tubes. J Exp Bot 58: 65–74

6. Carol RJ, Dolan L (2002) Building a hair: tip growth in *Arabidopsis thaliana* root hairs. Phil Trans R Soc Lond B 357: 815–821

7. Carol RJ, Takeda S, Linstead P, Durrant MC, Kakesova H, Derbyshire P, Drea S, Zarsky V, Dolan L (2005) A RhoGDP dissociation inhibitor spatially regulates growth in root hair cells. Nature 438: 1013–1016

8. Chapman JM, Muhlemann JK, Gayomba SR, Muday GK (2019) RBOH-dependent ROS synthesis and ROS scavenging by plant specialized metabolites to modulate plant development and stress responses. Chem Res Toxicol 32: 370–396

9. Clouse SD (2002) Arabidopsis mutants reveal multiple roles for sterols in plant development. Plant Cell 14: 1995–2000

10. Cole RA, Fowler JE (2006) Polarized growth: maintaining focus on the tip. Curr Opin Plant Biol 9: 579–588

11. Contento AL, Bassham DC (2012) Structure and function of endosomes in plant cells. J Cell Sci 125: 3511–3518

12. Costes SV, Daelemans D, Cho EH, Dobbin Z, Pavlakis G, Lockett S (2004) Automatic and quantitative measurement of protein-protein colocalization in live cells. Biophys J 86: 3993–4003

13. Denninger P, Reichelt A, Schmidt VAF, Mehlhorn DG, Asseck LY, Stanley CE, Keinath NF, Evers JF, Grefen C, Grossmann G (2019) Distinct RopGEFs successively drive polarization and outgrowth of root hairs. Curr Biol 29: 1854–1865

14. Dolan L, Duckett CM, Grierson C, Linstead P, Schneider K, Lawson E, Dean C, Poethig S, Roberts K (1994) Clonal relationships and cell patterning in the root epidermis of Arabidopsis. Development 120: 2465–2474

15. Foreman J, Demidchik V, Bothwell JH, Mylona P, Miedema H, Torres MA, Linstead P, Costa S, Brownlee C, Jones JD, et al (2003) Reactive oxygen species produced by NADPH oxidase regulate plant cell growth. Nature 422: 442–446

16. Geldner N, Dénervaud-Tendon V, Hyman DL, Mayer U, Stierhof YD, Chory J (2009) Rapid, combinatorial analysis of membrane compartments in intact plants with a multicolor marker set. Plant J 59: 169–178

17. Grierson C, Schiefelbein J (2002) Root hairs, The Arabidopsis book, 1: e0060. doi: 10.1199/tab.0060

18. Gillooly DJ, Simonsen A, Stenmark H (2001) Cellular functions of phosphatidylinositol3-phosphate and FYVE domain proteins. Biochem J 355: 249–258

19. Grebe M, Xu J, Möbius W, Ueda T, Nakano A, Geuze HJ, Rook MB, Scheres B (2003) Arabidopsis sterol endocytosis involves actin-mediated trafficking via ARA6-positive early endosomes. Curr Biol 13: 1378–1387

20. Haas TJ, Sliwinski MK, Martínez DE, Preuss M, Ebine K, Ueda T, Nielsen E, Odorizzi G, Otegui MS (2007) The Arabidopsis AAA ATPase SKD1 is involved in multi vesicular endosome function and interacts with its positive regulator LYST-INTERACTING PROTEIN 5. Plant Cell 19: 1295–1312

21. Hao H, Fan L, Chen T, Li R, Li X, He Q, Botella MA, Lin J (2014) Clathrin and membrane microdomains cooperatively regulate RbohD dynamics and activity in Arabidopsis. Plant Cell 26: 1729–1745

22. Jones MA, Shen JJ, Fu Y, Li H, Yang Z, Grierson CS (2002) The Arabidopsis Rop2 GTPase is a positive regulator of both root hair initiation and tip growth. Plant Cell 14: 763– 776

23. Jones MA, Raymond MJ, Smirnoff N (2006) Analysis of the root-hair morphogenesis transcriptome reveals the molecular identity of six genes with roles in root-hair development in Arabidopsis. Plant J 45: 83–100

24. Jones MA, Raymond MJ, Yang Z, Smirnoff N (2007) NADPH oxidase-dependent reactive oxygen species formation required for root hair growth depends on ROP GTPase. J Exp Bot 58: 1261–1270

25. Kaya H, Takeda S, Kobayashi MJ, Kimura S, Iizuka A, Imai A, Hishinuma H, Kawarazaki T, Mori K, Yamamoto Y, et al (2019) Comparative analysis of the reactive oxygen species-producing enzymatic activity of Arabidopsis NADPH oxidases. Plant J 98: 291–300

26. Kiegle E, Gilliham M, Haselof, J, Tester M (2000) Hyperpolarisation-activated calcium channels found only in cells from the elongation zone of *Arabidopsis thaliana* roots. Plant J 21: 225–229

27. Lindsey K, Pullen ML, Topping JF (2003) Importance of plant sterols in pattern formation and hormone signalling. Trends Plant Sci 8: 521–525

28. Liu P, Zhang L, Li R, Wang Q, Niehaus K, Balusška F, Šamaj J, Lin J (2009) Lipid microdomain polarization is required for NADPH oxidase-dependent ROS signaling in *Picea meyeri* pollen tube tip growth. Plant J 60: 303–313

29. Mangano S, Denita-Juarez SP, Choi H-S, Marzol E, Hwang Y, Ranocha P, Velasquez SM, Borassi C, Barberini ML, Aptekmann AA, et al (2017) Molecular link between auxin and ROS-mediated polar growth. Proc Natl Acad Sci USA 114: 5289–5294

30. Menon AK (2002) Lipid transport - An overview. Semin Cell Dev Biol 13: 159–162

31. Miedema H, Bothwell JHF, Brownlee C, Davies JM (2001) Calcium uptake by plant cells—channels and pumps acting in concert. Trends Plant Sci 11: 514–519

32. Milhaud J, Hartmann MA, Bolard J (1988) Interaction of the polyene antibiotic filipin with model and natural membranes containing plant sterols. Biochim Biophys Acta 943: 315–325

33. Molendijk AJ, Bischoff F, Rajendrakumar CS, Friml J, Braun M, Gilroy S, Palme K (2001) *Arabidopsis thaliana* Rop GTPases are localized to tips of root hairs and control polar growth. EMBO J 20: 2779–2788

34. Monshausen GB, Bibikova TN, Messerli MA, Shi C, Gilroy S (2007) Oscillations in extracellular pH and reactive oxygen species modulate tip growth of Arabidopsis root hairs. Proc Natl Acad Sci USA 104: 20996–21001

35. Morel J, Clavero S, Mongrand S, Furt F, Fromentin J, Bessoule_J-J, Blein J-P, Simon-Plas F (2006) Proteomics of plant detergent-resistant membranes. Mol Cell Proteomics 5: 1396–1411

36. Mukherjee S, Maxfield FR (2000) Role of membrane organization and membrane domains in endocytic lipid trafficking. Traffic 1: 203–211

37. Nichols BJ, Kenworthy AK, Polishchuk RS, Lodge R, Roberts TH, Hirschberg K, Phair RD, Lippincott-Schwartz J (2001) Rapid cycling of lipid raft markers between the cell surface and Golgi complex. J Cell Biol 153: 529–542

38. Ovečka M, Berson T, Beck M, Derksen J, Šamaj J, Baluška F, Lichtscheidl IK (2010) Structural sterols are involved in both the initiation and tip growth of root hairs in *Arabidopsis thaliana*. Plant Cell 22: 2999–3019

39. Ovečka M, Lang I, Baluška F, Ismail A, Illéš P, Lichtscheidl IK (2005) Endocytosis and vesicle trafficking during tip growth of root hairs. Protoplasma 226: 39–54

40. Ovečka M, Lichtscheidl I, Šamaj J (2014) Live microscopy analysis of endosomes and vesicles in tip-growing root hairs. Methods Mol Biol 1209: 31–44

41. Ovečka M, Vaškebová L, Komis G, Luptovčiak I, Smertenko A, Šamaj J (2015) Preparation of plants for developmental and cellular imaging by light-sheet microscopy. Nat Protoc 10: 1234–1247

42a. Posé D, Castanedo I, Borsani O, Nieto B, Rosado A, Taconnat L, Ferrer A, Dolan L, Valpuesta V, Botella MA (2009) Identification of the Arabidopsis dry2/sqe1-5 mutant reveals a central role for sterols in drought tolerance and regulation of reactive oxygen species The Plant Journal 59: 63–76

42. Qi X, Zheng H (2013) Rab-A1c GTPase defines a population of the trans-Golgi network that is sensitive to endosidin1 during cytokinesis in Arabidopsis. Mol Plant 6: 847–859

43. Reyes FC, Buono R, Otegui MS (2011) Plant endosomal trafficking pathways. Curr Opin Plant Biol 14: 666–673

44. Sagi M, Fluhr R (2006) Production of reactive oxygen species by plant NADPH oxidases. Plant Physiol 141: 336–340

45. Sanderfoot AA, Kovaleva V, Bassham DC, Raikhel NV (2001) Interactions between syntaxins identify at least five SNARE complexes within the Golgi/prevacuolar system of the Arabidopsis cell. Mol Biol Cell 12: 3733–3743

46. Šamaj J, Müller J, Beck M, Böhm N, Menzel D (2006) Vesicular trafficking, cytoskeleton and signalling in root hairs and pollen tubes. Trends Plant Sci 11: 594–600

47. Sena F, Sotelo-Silveira M, Astrada S, Botella MA, Malacrida L, Borsani O (2017) Spectral phasor analysis reveals altered membrane order and function of root hair cells in Arabidopsis *dry2/sqe1-5* drought hypersensitive mutant. Plant Physiol Biochem 119: 224–231

48. Schiefelbein JW, Somerville C (1990) Genetic control of root hair development in *Arabidopsis thaliana*. Plant Cell 2: 235–243

49. Simons K, Toomre D (2000) Lipid rafts and signal transduction. Nat Rev Mol Cell Biol 1: 31–39

50. Sorek N, Gutman O, Bar E, Abu-Abied M, Feng X, Running MP, Lewinsohn E, Ori N, Sadot E, Henis YI, et al (2011) Differential effects of prenylation and S-acylation on type I and II ROPS membrane interaction and function. Plant Physiol 155: 706–720

51. Sorek N, Poraty L, Sternberg H, Buriakovsky E, Bar E, Lewinsohn E, Yalovsky S (2017) Activation status coupled transient S-acylation determines membrane partitioning of a plant Rho-related GTPase. Mol Cell Biol 37: 2144–2154

52. Srivastava V, Malm E, Sundqvist G, Bulone V (2013) Quantitative proteomics reveals that plasma membrane microdomains from poplar cell suspension cultures are enriched in markers of signal transduction, molecular transport, and callose biosynthesis. Mol Cell Proteomics 12: 3874–3885

53. Stanislas T, Hüser A, Barbosa IC, Kiefer CS, Brackmann K, Pietra S, Gustavsson A, Zourelidou M, Schwechheimer C, Grebe M (2015) Arabidopsis D6PK is a lipid domain-dependent mediator of root epidermal planar polarity. Nat Plants 1: 15162

54. Takeda S, Gapper C, Kaya H, Bell E, Kuchitsu K, Dolan L (2008) Local positive feedback regulation determines cell shape in root hair cells. Science 319: 1241–1244

55. Torres MA, Dangl JL, Jones JD (2002) Arabidopsis gp91phox homologues AtrbohD and AtrbohF are required for accumulation of reactive oxygen intermediates in the plant defence response. Proc Natl Acad Sci USA 99: 517–522

56. Ueda T, Uemura T, Sato MH, Nakano A (2004) Functional differentiation of endosomes in Arabidopsis cells. Plant J 40: 783–789

57. Uemura T, Ueda T, Ohniwa RL, Nakano A, Takeyasu K, Sato MH (2004) Systematic analysis of SNARE molecules in Arabidopsis: dissection of the post-Golgi network in plant cells. Cell Struct Func 29: 49–65

58. Véry A-A, Davies JM (2000) Hyperpolarization-activated calcium channels at the tip of Arabidopsis root hairs. Proc Natl Acad Sci USA 97: 9801–9806

59. Viotti C, Bubeck J, Stierhof YD, Krebs M, Langhans M, van den Berg W, van Dongen W, Richter S, Geldner N, Takano J, et al (2010) Endocytic and secretory traffic in Arabidopsis merge in the trans-Golgi network/early endosome, an independent and highly dynamic organelle. Plant Cell 22: 1344–1357

60. Voigt B, Timmers A, Šamaj J, Hlavacka A, Ueda T, Preuss M, Nielsen E, Mathur J, Emans N, Stenmark H, et al (2005) Actin-based motility of endosomes is linked to the polar tip growth of root hairs. Eur J Cell Biol 84: 609–621

61. von Wangenheim D, Rosero A, Komis G, Šamajová O, Ovečka M, Voigt B, Šamaj J (2016) Endosomal interactions during root hair growth. Front Plant Sci 6: 1262

62. Wachtler V, Rajagopalan S, Balasubramanian MK (2003). Sterol-rich plasma membrane domains in the fission yeast *Schizosaccharomyces pombe*. J Cell Sci 116: 867–874

63. Wymer CL, Bibikova TN, Gilroy S (1997) Cytoplasmic free calcium distributions during the development of root hairs of *Arabidopsis thaliana*. Plant J 12: 427–439

64. Zhou X, Xiang Y, Li C, Yu G (2020) Modulatory role of reactive oxygen species in root development in model plant of *Arabidopsis thaliana*. Front Plant Sci 11: 485932

